# mTOR regulates longevity through a bile-acid like hormonal mechanism and DHS-26/DHRS1

**DOI:** 10.64898/2026.05.13.724957

**Authors:** Klara Schilling, Alex Zaufel, Kaylee M. Morris, Anna Löhrke, Ratni Saini, Hans-Joachim Knölker, Tarek Moustafa, Adam Antebi

## Abstract

The mTOR pathway is a central regulator of cellular metabolism and growth whose downregulation extends life span across taxa. In *C. elegans,* mTOR acts cell non-autonomously to influence organismal longevity, yet underlying mechanisms remain elusive. Here, we show that deletion of the TORC1 regulator, *raga-1*/*RRAGA*, enhances production of the bile acid–like hormone, dafachronic acid (DA), and extends life span dependent on DA-hormone biosynthetic genes and DA-cognate nuclear hormone receptor DAF-12, a homolog of mammalian farnesoid X receptor (FXR). Through functional genomic screens, we identify the evolutionarily conserved short chain dehydrogenase DHS-26/DHRS1 as a previously uncharacterized downstream regulatory target and effector of the mTOR-steroid axis essential for organismal longevity. Worm DHS-26 is expressed prominently in the canal associated neurons, cells which are essential to growth and development, suggesting a neuroendocrine mechanism. Murine DHRS1 also exhibits regulation by mTOR signaling and nuclear receptor FXR suggesting that the mTOR-DHS-26/DHRS1 axis is evolutionarily conserved. These findings suggest that mTOR signaling systemically impacts metazoan longevity through the regulation of bile acid–like hormone availability and nuclear receptor signal transduction.

## Introduction

Mechanistic target of rapamycin (mTOR) is a central integrator of nutrient, growth, and energy status that couples physiological state to cellular anabolism and catabolism. Under favorable conditions of nutrient availability, positive growth signaling, and high energy status, mTOR promotes anabolic processes, including ribogenesis and protein synthesis (Holz, Ballif et al. 2005), lipid and cholesterol production (Porstmann, Santos et al. 2008), and nucleotide metabolism (Ben-Sahra, Hoxhaj et al. 2016). Conversely, under adverse physiologic conditions, reduced mTOR activity dampens these anabolic processes, and promotes autophagy and lysosome biogenesis (Kim, Kundu et al. 2011, Roczniak-Ferguson, Petit et al. 2012), enhancing turnover and limiting growth and cell division. Given this central coordinating role, dysregulation of the mTOR pathway, not surprisingly, leads to a loss of cellular homeostasis, and is implicated in major diseases such as cancer (Sato, Nakashima et al. 2010), neurodegeneration (Spilman, Podlutskaya et al. 2010), sarcopenia and diabetes (Khamzina, Veilleux et al. 2005). mTOR dysregulation also drives organismal aging, and remarkably, pharmacological inhibition by rapamycin extends life span in model organisms across taxa (Harrison, Strong et al. 2009, Bjedov, Toivonen et al. 2010, Robida-Stubbs, Glover-Cutter et al. 2012), highlighting its crucial function in health and disease.

Many elegant mechanistic studies on mTOR have focused on defining its complexes, localization, regulation and signaling states within the cell, predominantly *in vitro*. How mTOR operates in a physiological context is less well understood. In the nematode *C. elegans*, disruption of *raga-1*, a component of the TORC1 pathway, reduces mTOR activity and extends life span (Schreiber, Pierce-Shimomura et al. 2010). Interestingly, downregulation of *raga-1* within the nervous system is sufficient to extend life span of the entire organism (Smith, Lanjuin et al. 2023). This finding implies that a cell non-autonomous mechanism downstream of mTOR regulates longevity, yet the nature of this signaling mechanism remains elusive.

Bile acids are derivatives of cholesterol that contain various modifications of the sterol nucleus and carboxylic acid moieties on their sidechain. Their amphipathic nature enables them to function as emulsifying agents during food digestion and absorption (Russell 2003). Beyond this, they act as endocrine hormones that signal through nuclear hormone receptors (NHR), such as farnesoid X receptor (FXR) (Makishima, Okamoto et al. 1999) and vitamin D receptor (VDR) (Mizwicki, Keidel et al. 2004), as well as G protein receptors (Kawamata, Fujii et al. 2003) to regulate cholesterol, lipid and glucose metabolism, with profound effects on animal physiology. In *C. elegans*, bile acid-like steroids, termed the dafachronic acids (DAs), act through the nuclear hormone receptor DAF-12, a homolog of mammalian FXR and VDR, to regulate the dauer diapause, developmental timing, and adult longevity (Antebi, Yeh et al. 2000). In response to environmental stress and nutrient cues, the unliganded receptor promotes developmental arrest at the dauer stage (Ludewig, Kober-Eisermann et al. 2004), whereas in favorable conditions the liganded receptor promotes larval developmental progression as well as adult longevity in several contexts (Antebi, Yeh et al. 2000, Gerisch, Weitzel et al. 2001, Gerisch, Rottiers et al. 2007). While mTOR signaling also affects the dauer diapause and adult longevity (Jia, Chen et al. 2004), the relationship between mTOR and DA/DAF-12 signaling remains poorly understood.

Here, we hypothesized that DA signaling might be linked to mTOR enabling a cell non-autonomous mechanism governing longevity. Our study suggests that *raga-1* regulates sterol availability and DA production, and DA/DAF-12 work downstream of *raga-1* to mediate life span extension. A functional genomic screen identifies an evolutionarily conserved, but uncharacterized short chain dehydrogenase *dhs-26*/*DHRS1*, as a key output of the mTOR-steroid axis mediating longevity. These findings imply that mTOR may work systemically via bile acid-like steroids and NHR transduction to regulate longevity in metazoans.

## Results

### Steroid signaling is required for longevity induced by decreased mTOR signaling

To test the idea that mTOR and DA/DAF-12 signaling interact to influence longevity, we used mutants deficient in the first and last steps of DA biosynthesis *daf-36(k114)/*Rieske oxygenase (Wollam, Magomedova et al. 2011) and *daf-9(rh50)/CYP27A1* (Gerisch, Weitzel et al. 2001) (Fig. 1a), as well as the NHR *daf-12(rh61rh411)/*FXR in the background of *raga-1(ok701),* a genetic model for reduced TORC1 signaling. These alleles are presumptive nulls, with the exception of *daf-9(rh50),* a reduction-of-function mutation.

**Fig. 1.**
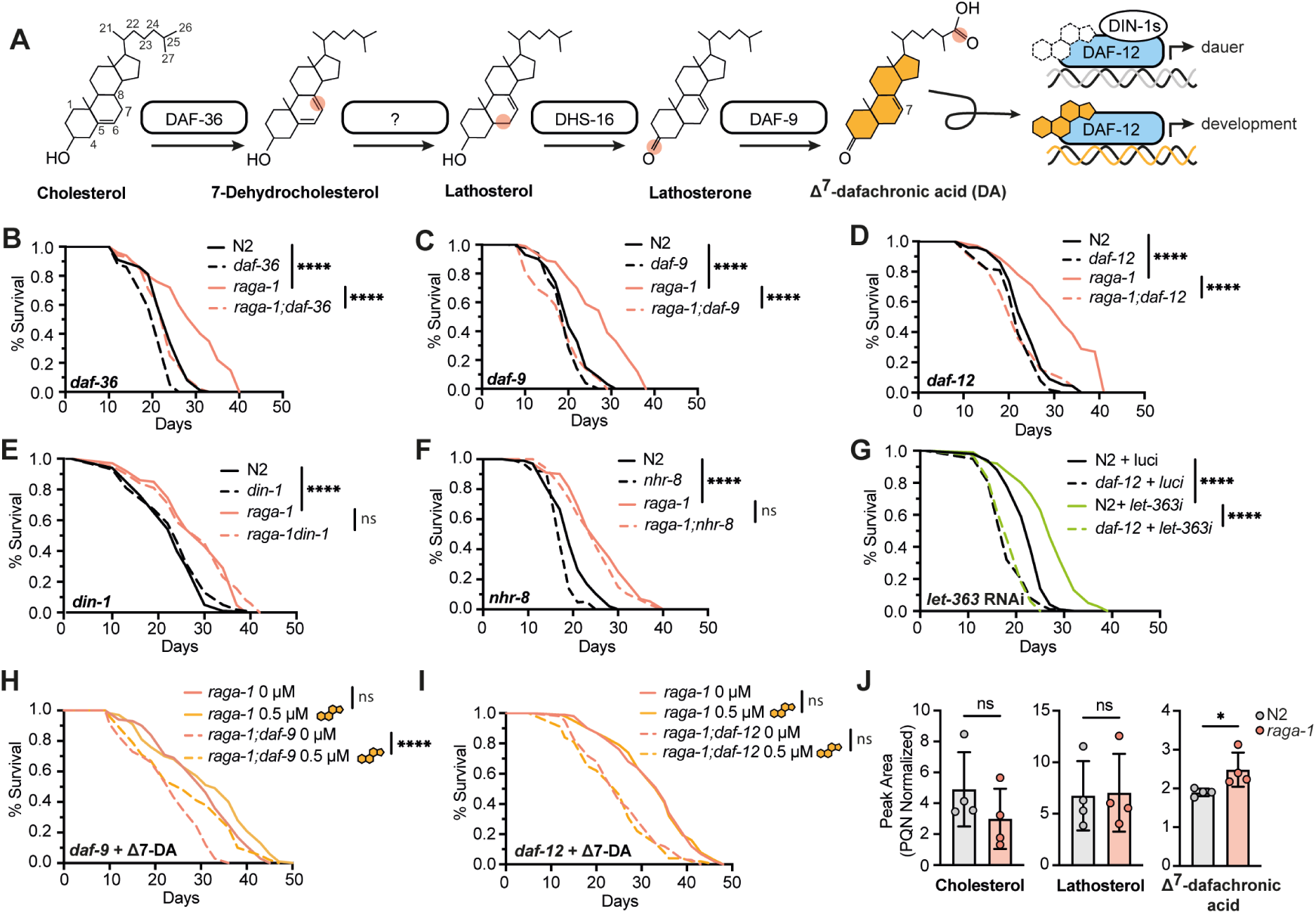
Steroid signaling is required for longevity induced by decreased mTOR signaling. **a**, Schematic overview of the endogenous dafachronic acid biosynthesis pathway in *C. elegans*. In the first step, the Rieske oxygenase DAF-36/Neverland serves as a cholesterol 7-desaturase, catalyzing the oxidation of cholesterol to 7-dehydrocholesterol (Wollam, Magomedova et al. 2011). An unknown enzyme catalyzes the reduction of the Δ-5 bond to lathosterol. Short-chain dehydrogenase DHS-16 carries out the oxidation of the 3-hydroxy to 3-keto derivative to yield lathosterone (Wollam, Magner et al. 2012). In the final step, cytochrome P450 DAF-9/CYP27A1 catalyzes the oxidation of the sterol sidechain to the carboxylic acid moiety (Motola, Cummins et al. 2006), resulting in the formation of Δ7-DA. Δ7-DA functions as a ligand for the nuclear hormone receptor DAF-12/FXR to promote development. Unliganded DAF-12 binds to its corepressor DIN-1s inducing dauer diapause. **b-f**, Representative life spans comparing wild type (N2), *raga-1(ok701),* and steroid mutants. b) *daf-36(k114)* and *raga-1;daf-36*, c) *daf-9(rh50)* and *raga-1;daf-9*, d) *daf-12(rh61rh411)* and *raga-1;daf-12*, e) *din-1(dh127)* and *raga-1;din-1* f) *nhr-8(hd117)* and *raga-1;nhr-8*. Statistical analysis for life spans was performed using the log-rank (Mantel–Cox) test. ns, non-significant (p>0.05), **** (p<0.001). Data and statistics are summarized in Supplementary Table 1. **g,** Representative life spans comparing N2 and *daf-12(rh61rh411)* on *let-363*/mTOR and control (*luc*) RNAi. Eggs were cultivated on regular NGM plates until day 1 of adulthood, then worms were transferred to the respective RNAi plates. Statistical analysis for life spans was performed using the log-rank (Mantel-Cox) test. **** (p<0.005). Data and statistics are summarized in Supplementary Table 1. **h-i**, Life span upon 0.5 µM Δ7-DA supplementation or vehicle (EtOH) control egg-on, comparing h) *raga-1(ok701)* and *raga-1;daf-9(rh50),* and i) *raga-1(ok701)* and *raga-1;daf-12*. Statistical analysis for life spans was performed using the log-rank (Mantel–Cox) test. ns (p>0.05), *** (p<0.005), n=2. Data and statistics are summarized in Supplementary Table 1. **j,** Relative endogenous concentrations of cholesterol, lathosterol, and Δ7-dafachronic acid (DA) from N2 and *raga-1(ok701)* day 1 adults. DA concentrations were normalized to PQN, n=4. Mean ± SD. Statistical analysis was performed using the Student’s t-test. ns (p>0.05), * (p<0.01)

As expected, *raga-1* single mutants were long-lived, with a 20-30% increase in life span (Fig. 1b-f) as described previously (Schreiber, Pierce-Shimomura et al. 2010). Consistent with our hypothesis, mutation of *daf-36, daf-9,* or *daf-12* abolished *raga-1* mutation-induced longevity (Fig. 1b-d). By contrast, loss of *din-1*/SHARP, a co-repressor that works together with unliganded *daf-12*, or *nhr-8*/LXR, a NHR closely related to *daf-12*, had no effect (Fig. 1e-f). In accord with a hormonal mechanism, supplementation with Δ7-DA increased life span in DA deficient *raga-1;daf-9* (Fig. 1h) but not in the receptor deficient *raga-1;daf-12* animals (Fig. 1i). Notably, *daf-12* mutation also abolished the life span extension of *let-363*/TOR RNAi knockdown (Fig. 1g) showing pathway-rather than gene-specific interaction. Altogether, these findings support the idea that DA acts through its canonical nuclear receptor *daf-12* to promote life span extension arising from reduced mTOR signaling.

Since DA/DAF-12 acts genetically downstream of mTOR signaling, we asked whether mTOR modulates DA pathway signaling itself. To test this idea, we directly measured Δ7-DA hormone levels by gas chromatography-mass spectrometry. Interestingly, Δ7-DA levels, but not cholesterol and lathosterol, were modestly but significantly upregulated by 20% in *raga-1* mutant compared to N2, suggesting that mTOR inhibition promotes DA signaling via Δ7-DA (Fig. 1j).

We next wondered whether *daf-12*/DA pathway might in turn affect mTOR signaling. To assess mTOR activity we used two established methods in *C. elegans*, HLH-30/TFEB nuclear localization and AMPK phosphorylation (Lapierre, De Magalhaes Filho et al. 2013, Zhang, Lanjuin et al. 2019). As expected, *raga-1* loss resulted in enhanced nuclear localization of *hlh-30*/TFEB, while loss of *daf-36* had no consistent effect, either in wild type or *raga-1* backgrounds (Supplementary Fig. 1a-b). Similarly, *raga-1* mutation enhanced AMPK phosphorylation, while mutations in DA/DAF-12 pathway components showed no significant effect (Supplementary Fig. 1c-g). These findings indicate that DA/DAF-12 signaling has little or no impact on mTOR signaling, working downstream of the pathway.

### *daf-12* mutation modulates the *raga-1* mutant transcriptome

To gain insight into links between mTOR and steroid signaling and possible downstream mechanisms, we conducted transcriptomic analyses, comparing wild type, *raga-1*, *daf-12,* and *raga-1;daf-12* mutants at day 1 of adulthood. Principle Component Analysis (PCA) showed that *raga-1;daf-12* mutants separate from *raga-1* along PC1, while mutation of *raga-1* compared to wild type separates along PC2 (Fig. 2a). Additionally, PC1 likely separates differences in longevity. Volcano plots comparing *raga-1* mutants and N2 transcriptomes showed markedly significant changes (Adj.p<0.05), with 4319 down-regulated differentially expressed genes (DEGs) and 4578 up-regulated DEGs (Supplementary Fig. 2a). Down-regulated genes were enriched in genes related to ribosome and nucleocytoplasmic transport (Supplementary Fig. 2b), while up-regulated genes showed enrichment in metabolic pathway genes, as well as lysosomal and peroxisomal genes (Supplementary Fig. 2c). Comparisons of *raga-1;daf-12* to *raga-1* mutants showed 970 down-regulated and 1314 up-regulated DEGs (Fig. 2b), revealing that *daf-12* strongly modulates the *raga-1* transcriptome. KEGG gene ontology analysis of down-regulated genes from *raga-1;daf-12/raga-1* animals comparisons showed enrichment for TCA cycle, one-carbon metabolism, fatty acid synthesis and degradation, amino acid metabolism, glutathione metabolism, and peroxisomal pathways, indicating a rewiring of metabolic pathways and organelle function (Fig. 2c). Up-regulated genes were enriched for Hippo signaling, lysosomal pathways, and axon regeneration. Altogether, these findings suggest dysregulated metabolism, growth signaling and cellular remodeling (Fig. 2c).

**Fig. 2.**
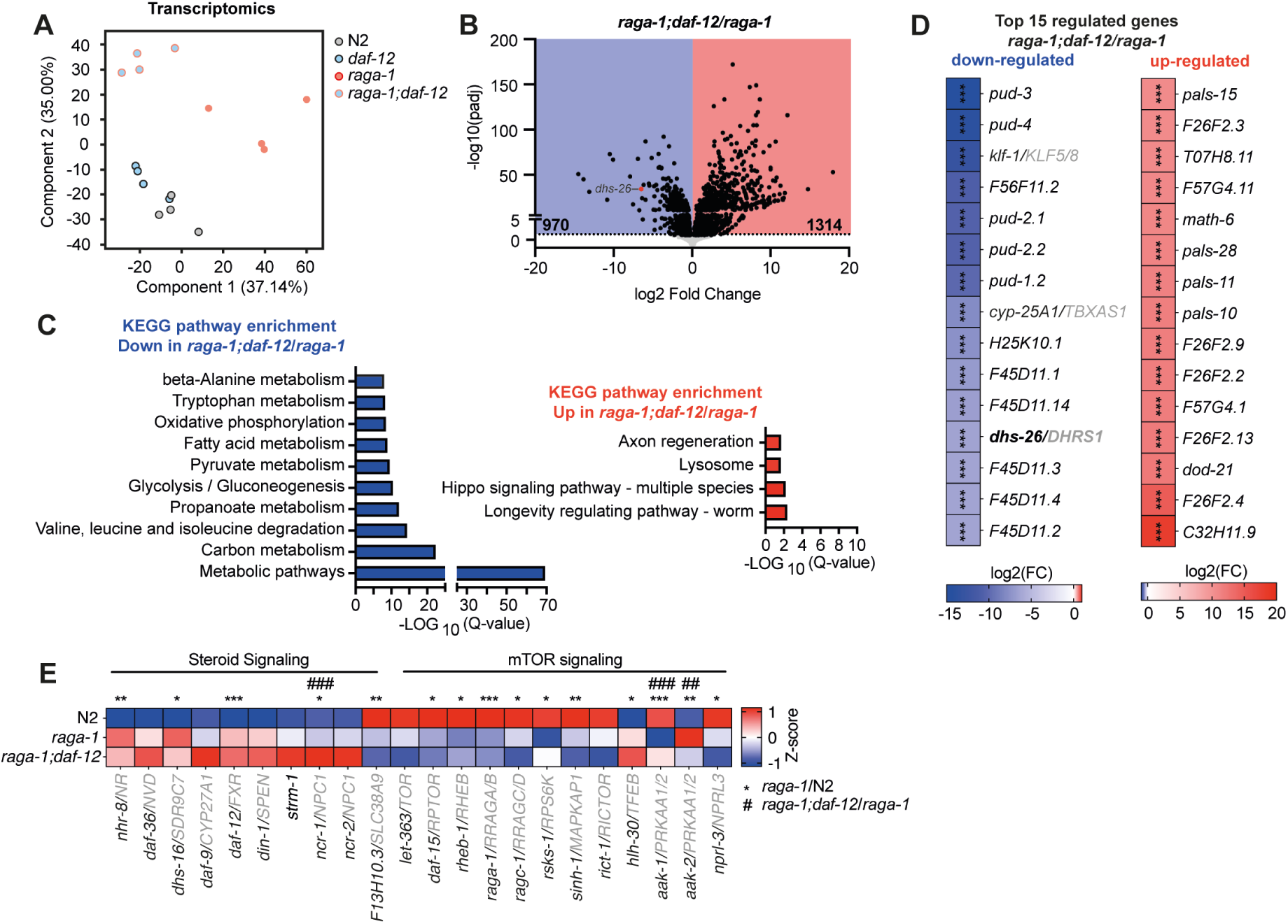
daf-12 modulates the raga-1 mutant transcriptome. **a**, Principal component analysis of wild type (N2), *raga-1(ok701), daf-12(rh61rh411),* and *raga-1;daf-12* mutant transcriptomes in day 1 adults. **b**, Volcano plot of differentially expressed genes comparing *raga-1;daf-12/raga-1(ok701).* Black dots mark significantly changed genes (Adj.p<0.05), grey dots mark non-significantly changed genes (Adj.p>0.05), *dhs-26* is highlighted as a red dot. Blue area highlights 970 significantly down-regulated genes; the red area highlights 1314 significantly up-regulated genes. **c**, KEGG pathway enrichment (FDR<0.05) of significantly (Adj.p<0.05) down-regulated (blue) and up-regulated genes (red) comparing *raga-1;daf-12/raga-1(ok701).* The gene list of KEGG enrichments is contained in Supplementary Table 3. **d**, Heatmap of log2(FC) of the top 15 down-regulated (blue) and top 15 up-regulated genes (red) comparing *raga-1;daf-12/raga-1(ok701).* *** (Adj.p<0.005). **e**, Z-score heatmap of genes involved in steroid signaling and mTOR signaling comparing N2, *raga-1(ok701)* and *raga-1;daf-12.* Comparing *raga-1*/N2, * (Adj.p<0.05), ** (Adj.p<0.01), *** (Adj.p<0.005). Comparing *raga-1;daf-12/raga-1(ok701)*, ## (Adj.p<0.01), ### (Adj.p<0.005).

A closer look into the top 15 up- and down-regulated genes ranked by fold-change revealed genes encoded in close genomic proximity, including families of *pud*, *F45D11, F26F2, F57G4*, and *pals* genes (Fig. 2d), consistent with previous observations that *daf-12* often regulates gene blocks throughout the genome (Shostak, Van Gilst et al. 2004). Additionally, single genes *klf-1*, *cyp-25A1*, and *dhs-26* were among the top 15 most down-regulated genes, suggesting they might be direct *daf-12* targets.

We also analyzed the differential gene expression of known steroid and mTOR pathway related genes in our transcriptomics dataset. Z-score analysis comparing the transcriptome of N2 and *raga-1* mutant animals indicated up-regulation of some steroid pathway genes (Fig. 2e), including biosynthetic gene *dhs-16* and nuclear hormone receptors *nhr-8* and *daf-12*. Further, lysosomal cholesterol transporter *ncr-1* was significantly up-regulated, while *SLC38A9*, which amongst other functions is responsible for cholesterol sensing (Castellano, Thelen et al. 2017), was significantly down-regulated (Fig. 2e). No significant change in the expression of the hormone biosynthesis genes *daf-36* and *daf-9* was observed at the transcriptional level (Fig. 2e). At the protein level, *raga-1* had no significant impact on expression of DA pathway components using *daf-36, daf-9* and *daf-12* translational fusion protein reporters (Supplementary Fig. 2d-k), confirming previous transcriptional results for *daf-36* and *daf-9,* while DHS-16::GFP reporter levels were decreased (Supplementary Fig. 2f-g).

Transcript levels of known mTOR pathway related genes did not reveal significant changes when comparing *raga-1* and *raga-1;daf-12* mutants. Exceptionally, AMPK subunits *aak-1* and *aak-2* showed significant regulation on mRNA level (Fig. 2e), but not protein level (Supplementary Fig. 5c). Unchanged expression of mTOR components is in line with the observed unchanged mTOR activity (Supplementary Fig. 1b-g).

Our data suggest that mTOR dependent regulation of steroid signaling might not be mediated through changes in transcription or translation, but potentially acts by regulating DA production and DAF-12 activity.

### Functional genomic screen reveals DAF-12 regulated gene *dhs-26* affects *raga-1* life span

To identify *daf-12* regulated genes mediating the observed effect on *raga-1* mutant life span, we performed a functional genomics RNAi screen looking for candidates that abolished longevity similar to *daf-12* mutation. For screen candidate selection, we first performed Pearson correlation between DEGs in *raga-1;daf-12*/*raga-1* and *raga-1*/N2 animals, resulting in 536 genes that were down-regulated in short-lived *raga-1;daf-12* but up-regulated in long-lived *raga-1* mutant animals (Fig. 3a). We then restricted candidates to a minimal log2 fold-change of 0.5, minimal read count of 10 per million, availability of a suitable RNAi clone as well as gene conservation in humans, mice, or *D. melanogaster* (Fig. 3b), leaving 185 genes. To test the selected candidate genes, we induced RNAi knockdown by feeding in *raga-1* mutants from egg-on and scored the extent of adult survival (Fig. 3b). As expected, most genes reduced *raga-1* mutant life span upon RNAi knockdown compared to *raga-1 luci* control, while a few genes further increased *raga-1* mutant life span (Fig. 3c). As observed previously, *raga-1;daf-12 luci* control showed reduced life span compared to *raga-1 luci* (Fig. 3c). Focusing on genes that showed a highly significant change in *raga-1* mutant life span upon knockdown (p<0.001) resulted in 29 genes (Fig. 3d). Among them, only the dehydrogenase *dhs-26*/*DHRS1* emerged as both significant for life span regulation as well as a robustly regulated in the transcriptome, as mentioned above (Fig. 2d).

**Fig. 3.**
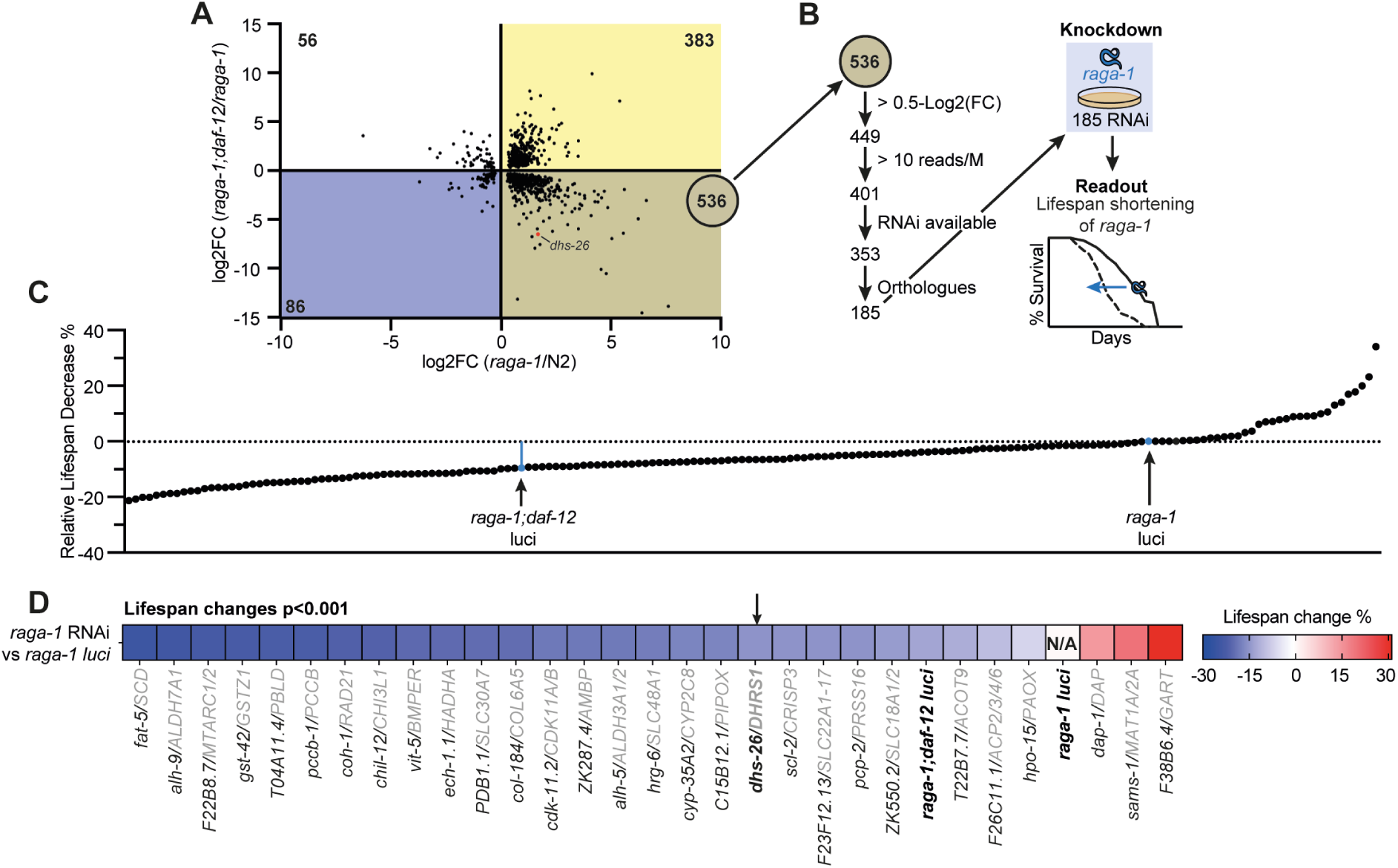
Functional genomic screen reveals that DAF-12 regulated gene dhs-26 affects raga-1 life span. **a**, Pearson correlation analysis of the Log2(FC) of all genes significantly changed in expression comparing *raga-1;daf-12/raga-1(ok701)* (Y axis) and *raga-1(ok701)*/N2 (X axis). 536 genes are significantly down-regulated in *raga-1;daf-12/raga-1(ok701)* and up-regulated in *raga-1(ok701)*/N2 (brown). *dhs-26* gene is indicated by the red dot. 383 genes are significantly up-regulated in *raga-1;daf-12/raga-1(ok701)* and *raga-1(ok701)*/N2 (yellow). 86 genes are significantly down-regulated in *raga-1;daf-12/raga-1(ok701)* and *raga-1(ok701)/*N2 (blue). 56 genes are significantly up-regulated in *raga-1;daf-12/raga-1(ok701)* and down-regulated in *raga-1(ok701)/*N2 (white). **b,** The 536 candidate genes shown were filtered using Log2(FC)>0.5, reads/million>10, availability of the respective RNAi clone, presence of gene orthologues in human, mice or *D. melanogaster*, resulting in 185 candidate genes. Schematic overview of the functional genomic screen. 185 genes were individually knocked down by RNAi in *raga-1* mutants to screen for influence on adult life span. **c**, Bar graph displaying the relative mean life span change (%) of *raga-1* mutants fed RNAi bacteria of candidate genes. Mean life span was assessed by feeding worms with RNAi bacteria egg-on and scoring for survival every two to three days. *raga-1(ok701)* and *raga-1;daf-12* grown on luciferase (*luc*) RNAi were included as reference controls. Statistical analysis was performed using the log-rank (Mantel-Cox) test, n=1. **d**, Heat map displaying significant life span change (p<0.001) induced by RNAi clones on *raga-1* life span displayed in c.

DHS-26 is an NADPH dependent short chain dehydrogenase orthologous to mammalian DHRS1. It exhibits 37% identity and contains similar domains and overall predicted 3D structure to its human counterpart (r.m.s.d. 2.748 Å) (Fig. 4a-c, Supplementary Fig. 3a-b). Human DHRS1 has been shown to biochemically metabolize several steroids, xenobiotics and other small molecules *in vitro* (Zemanova, Navratilova et al. 2019), and is highly expressed in liver and adrenals, and at lower levels in heart, skeletal muscle, pancreas, kidney and brain (Consortium 2013). However, its endogenous physiological function is completely unknown. We therefore sought to further characterize *dhs-26* in *C. elegans*.

**Fig. 4.**
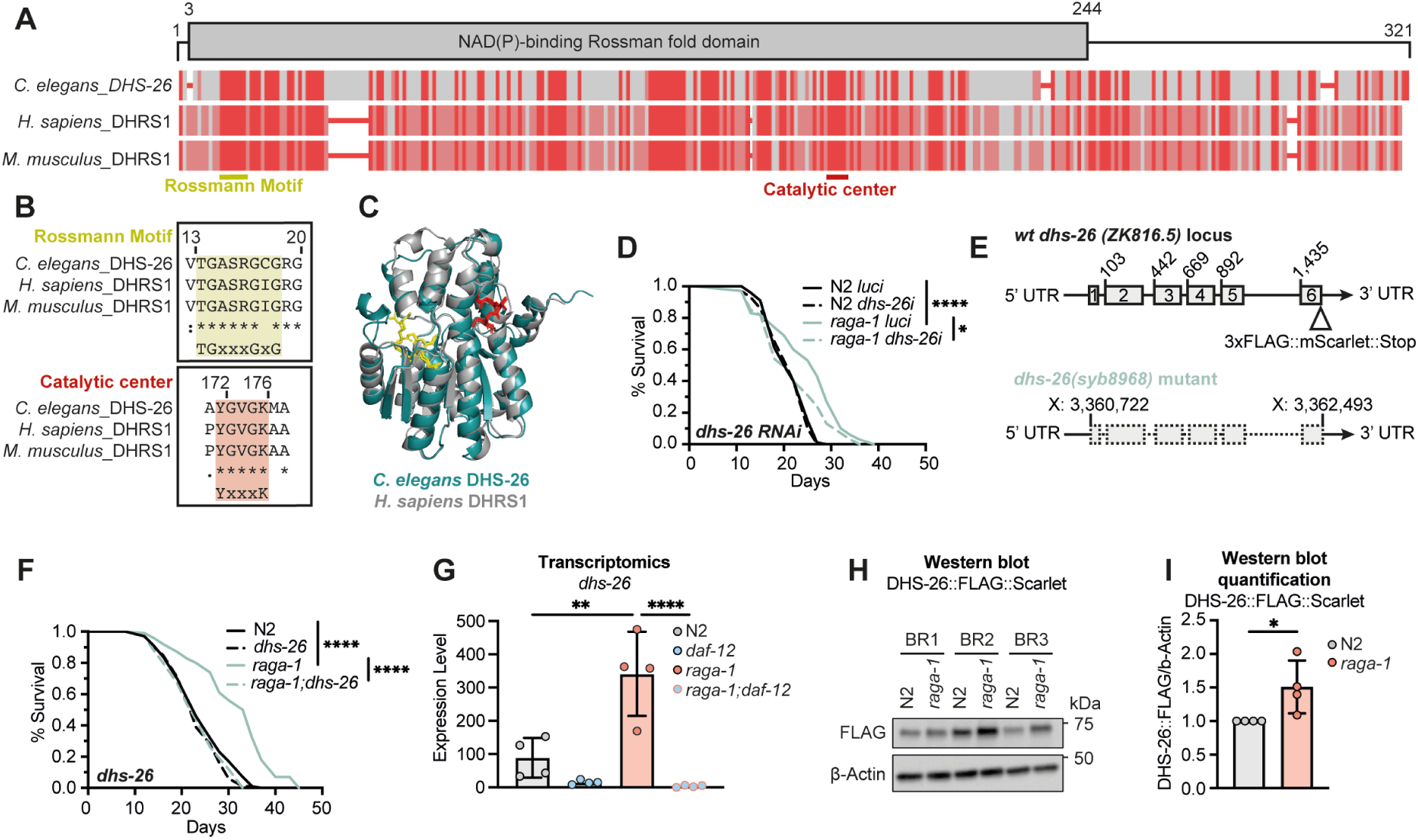
raga-1 mutants require the highly conserved dehydrogenase DHS-26/DHRS1 for life span extension. **a**, Multiple sequence amino acid alignment of DHS-26 with orthologs from other species using constraint-based multiple sequence alignment tool (COBALT). Red=identical amino acids, grey=non-identical. Above, the NAD(P)-binding Rossman fold domain is shown. Below, the Rossman motif (yellow) and the catalytic center (red) are highlighted. **b**, Sequence alignment of the Rossman motif (TGxxxGxG, yellow), and the catalytic center (YxxxK, red) of different organisms. **c**, Structural alignment of the AlphaFold 3-predicted model of human DHRS1 (PDB Q96LJ7, grey) with the AlphaFold 3-predicted model of *C. elegans* DHS-26 (PDB Q23612, turquoise). The Rossman motif (yellow) and catalytic center (red) are highlighted. r.m.s.d. = 2.748 Å. **d,** Representative life span comparing wild type (N2) and *raga-1(ok701)* on *dhs-26/DHRS1* and control (*luc*) RNAi treatment egg-on. Statistical analysis was performed using the log-rank (Mantel-Cox) test. * (p<0.05), **** (p<0.001), n=3. Data and statistics are summarized in supplementary Table 1. **e**, Schematic of the genomic *dhs-26* locus and structure of the *dhs-26(syb8968)* deletion mutant (X: 3,360,722-3,362,493). Triangle indicates the endogenous insertion site of *flag::Scarlet* tag at the C-terminus. **f**, Representative life span analysis of N2, *dhs-26(syb8968), raga-1(ok701),* and *raga-1;dhs-26* mutants. Statistical analysis was performed using the log-rank (Mantel-Cox) test. **** (p<0.001), n=4. All data and statistics are summarized in supplementary Table 1. **g,** *dhs-26* mRNA levels measured by RNA sequencing in the indicated mutant genotypes. Statistical analysis was performed using One-way ANOVA, ** (p<0.01), **** (p<0.001), n=4. **h**, Immunoblots with lysates from 500 day 1 N2 and *raga-1(ok701)* worms with endogenously tagged DHS-26::FLAG::Scarlet, probed with the indicated antibodies. DHS-26::FLAG::Scarlet fusion protein is expected at 73 kDa. n=4. **i,** Quantification of immunoblots displaying fold change of FLAG-band intensities normalized to β-Actin loading control and N2. Statistical analysis was performed using Student’s t-test, * (p<0.05), n=4.

We first CRISPR engineered a full knockout mutation at the endogenous locus, *dhs-26(syb8968)*, lacking all six exons (Fig. 4e). The deletion mutant was viable and had normal body size, and brood size comparable to wild type (Supplementary Fig. 3c-e). We next ascertained its influence on longevity to validate that the deletion exhibits the same life span effects as RNAi knockdown (Fig. 4d-f). On its own, deletion of *dhs-26* had little or no effect on life span, living to a similar extent as wild type. However, *dhs-26* deletion completely abolished *raga-1* mutant longevity in the *raga-1;dhs-26* double mutant (Fig. 4f), confirming the RNAi results from our screen. Of note, *raga-1;dhs-26* animals showed levels of life span suppression reminiscent of steroid gene and *daf-12* mutants (Fig. 1b-d).

### Regulatory crosstalk between DHS-26 and steroid signaling

We next sought to investigate *dhs-26* expression and regulation. Levels of *dhs-26* mRNA as measured by RNA-seq were elevated in *raga-1* mutants compared to N2 and reduced in the *raga-1;daf-12* background (Fig. 4g), which we validated using RT qPCR (Supplementary Fig. 3f). To monitor protein expression, we CRISPR engineered an endogenously C-terminally tagged DHS-26::FLAG::Scarlet reporter construct (Fig. 4e). Western blotting confirmed that DHS-26 protein was upregulated in *raga-1* mutants (Fig. 4i). Fluorescence microscopy of worms *in vivo* revealed that DHS-26::FLAG::Scarlet (Supplementary Fig. 3g) was expressed prominently in the canal associated neuron (CAN) (Supplementary Fig. 3g), a cell that extends processes along the excretory canal, essential for growth and development (Forrester and Garriga 1997, Chien, Wolf et al. 2019). We also observed occasional weak DHS-26 expression in a handful of unidentified head neurons and glial cells under DA supplementation (Supplementary Fig. 3g). To validate the functionality of the DHS-26::FLAG::Scarlet construct, the body morphology of *raga-1;dhs-26::FLAG::Scarlet* reporter animals was compared with *raga-1;dhs-26* double mutants which exhibited increased curling, referred to as the omega phenotype (Supplementary Fig. 3h-i). *raga-1;dhs-26::FLAG::Scarlet* animals showed no increase in omega phenotype occurrence compared to *raga-1* (Supplementary Fig. 3i), showing that the DHS-26 fusion protein is functional.

Notably, the dependence of *dhs-26* mRNA expression on *daf-12* shown above (Fig. 4g) suggested DA via DAF-12 could regulate *dhs-26*. To test this idea, we assessed DHS-26::Scarlet expression in *daf-36* mutants, deficient in DA production, as well as in nuclear hormone receptor *daf-12* mutants, focusing on the CAN. Consistent with a hormonal mechanism, loss of DA hormone or hormone receptor reduced the expression of *dhs-26* (Fig. 5a-b), while Δ7-DA supplementation robustly restored expression to *daf-36* but not *daf-12* mutants. As an additional test, we assessed reporter expression in wild type animals under exogenous cholesterol deprivation, which also lowers DA levels. Again, DHS-26 expression was reduced (Fig. 4c-d), in accord with the idea that *dhs-26* is regulated by DA/DAF-12 signaling. Whether regulation is direct or indirect is not clear. Existent DAF-12 chromatin immunoprecipitation data do not explicitly identify binding at the *dhs-26* locus (Hochbaum, Zhang et al. 2011), despite DAF-12 expression in the CAN (Supplementary Fig. 3g), and DAF-12 binding motifs (Ao, Gaudet et al. 2004, Shostak, Van Gilst et al. 2004) being present in the *dhs-26* promoter region. The absence of a robust signal might arise from low levels of DAF-12, alongside low *dhs-26* expression in only a handful of cells.

**Fig. 5.**
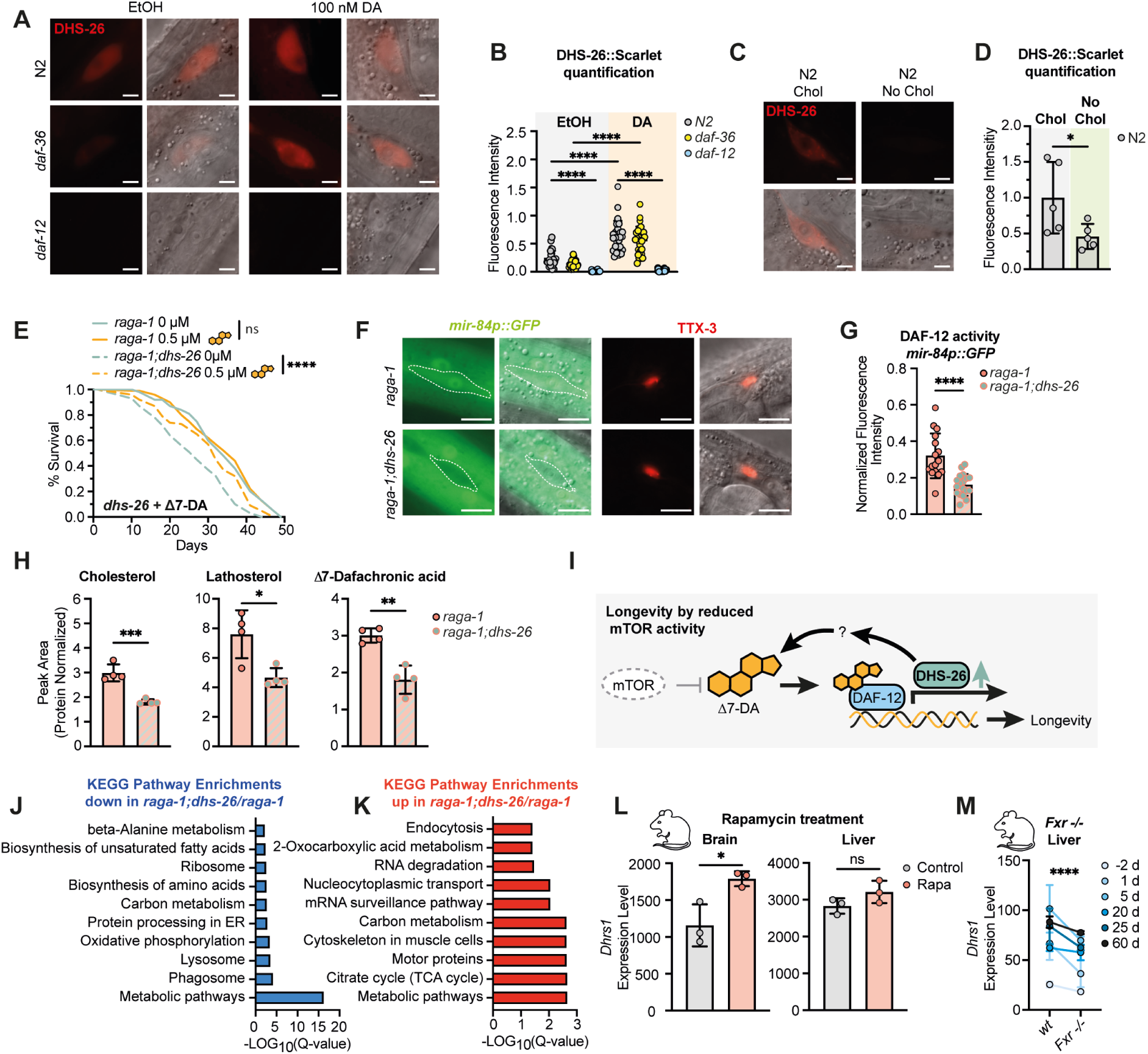
DHS-26/DHRS1 regulates steroid signaling and is required for life span extension induced by reduced mTOR. **a**, Representative images of DHS-26::FLAG::Scarlet (DHS-26::Scarlet) expression in the canal associated neurons (CAN) of day 1 adults, lateral aspect midbody. Worms were supplemented with 100 nM Δ7-DA or vehicle control (EtOH) egg-on. Scale bar, 5 µm. **b**, Representative quantification of DHS-26::Scarlet fluorescence intensity from a. Fluorescence intensity of each region of interest (ROI) was calculated by subtracting the background fluorescence of each image from the ROI. Statistical analysis was performed using One-way ANOVA, **** (p<0.001), n=3. **c**, Representative images of DHS-26::Scarlet expression in the CAN of day 1 N2 adults, lateral aspect midbody. Worms were cultured on regular NGM plates containing 5 µg/mL cholesterol (Chol) or on NGM plates without cholesterol supplementation (No Chol) egg-on. n=5, Scale bar, 5 µm. **d**, Quantification of DHS-26::Scarlet fluorescence intensity from c. Fluorescence intensity of each region of interest (ROI) was calculated by subtracting the background fluorescence of each image from the ROI. Statistical analysis was performed using paired t-test, * (p<0.005), n=5. **e**, Representative life span of *raga-1(ok701)* and *raga-1;dhs-26* animals supplemented with 0.5 µM Δ7-DA or EtOH control. Statistical analysis was performed using the log-rank (Mantel-Cox) test. ns (p>0.05), **** (p<0.001), n=3. Data and statistics are summarized in supplementary Table 1. **f,** Representative images of *mir-84p::GFP* expression in seam cell V2 of animals (highlighted) at larval stages L4.3-L4.5 (Mok, Sternberg et al. 2015). Larval stage was assessed by vulval morphology. Differential interference contrast (DIC) and fluorescence images were taken at midbody for *mir-84p::GFP*, and head for TTX-3::RFP of the same worm. Scale bar, 10 µm. **g**, Representative quantification of *mir-84p::GFP* expression in seam cell V2 of *raga-1(ok701)* and *raga-1;dhs-26* animals. Fluorescence intensity of each region of interest (ROI) was calculated by subtracting the background fluorescence of each image from the ROI, and was additionally normalized to TTX-3::RFP expressed in AIY interneurons. Statistical analysis was performed using Student’s t-test, **** (p<0.001), n=3. **h**, Relative concentrations of Δ7-dafachronic acid (DA) and its precursors cholesterol and lathosterol extracted from *raga-1(ok701)* and *raga-1;dhs-26* day 1 adults. Measured concentrations were normalized to protein levels. Statistical analysis was performed using Student’s t-test, * (p<0.05), ** (p<0.01), *** (p<0.005), n=4. **i**, Working model. mTOR inhibits DA biosynthesis. Reduced mTOR activity elevates DA levels. Increased DA binding to the nuclear hormone receptor DAF-12 drives the expression of DAF-12 target genes such as *dhs-26*, which positively regulates DA production. Activation of DAF-12 and DHS-26 additionally induce downstream processes linked to longevity. **j**, KEGG pathway enrichment (FDR<0.05) of significantly (Adj.p<0.05) downregulated (blue) genes comparing *raga-1;dhs-26/raga-1.*The gene list of KEGG enrichments is listed in supplemenrary Table 3. **k**, KEGG pathway enrichment (FDR<0.05) of significantly (Adj.p<0.05) upregulated genes (red) comparing *raga-1;dhs-26/raga-1(ok701).* The gene list of KEGG enrichments is listed in supplementary Table 3. **l***, Dhrs1* mRNA levels measured by RNA sequencing in the brain and liver of 15 day old female TK2KI animals treated with rapamycin or vehicle control (EtOH) by Siegmund *et al*. (Siegmund, Yang et al. 2017). Rapamycin and EtOH were administered in drinking water. During pregnancy the administered rapamycin concentration was 8 µg/ml, and 40 µg/ml after delivery. Untreated control litters were given water with equivalent concentrations (0.1%) of vehicle alone (100% ethanol). Statistical analysis was performed using Student’s t-test, ns (p>0.05), * (p<0.05), n=3. **m**, *DHRS1* mRNA levels measured by RNA sequencing in the liver of male C57BL/6 control and Fxr−/− mice by Peng *et al*. (Peng, Piekos et al. 2017). Animals were sacrificed at day −2 (gestational day 17.5), day 1 (exactly 24 hours after birth), and days 5, 20, 25, and 60 (collected at approximately 9:00 AM). Statistical analysis was performed using 2-Way ANOVA of paired mean difference, **** (p<0.0001), n=3.

As shown above, *dhs-26* is required for *raga-1* induced longevity similar to DA deficient mutants. Further, its human ortholog DHRS1 reportedly metabolizes steroids. We therefore postulated that *dhs-26* could play a role in steroid uptake, metabolism or DA production itself. To test this hypothesis, we first supplemented Δ7-DA to various genotypes and measured the impact on life span. We found that while Δ7-DA exposure had no effect on long-lived *raga-1* single mutants, it restored longevity to short lived *raga-1;dhs-26* double mutants (Fig. 5e), consistent with idea that *raga-1*;*dhs-26* animals are DA-deficient.

To further explore this notion, we used an established DA/DAF-12 dependent target gene, the microRNA *mir-84* (Bethke, Fielenbach et al. 2009), as a readout of DA/DAF-12 activity *in vivo*. In particular, we measured *mir-84p::gfp* promoter fusion expression in hypodermal seam cells, and normalized to a TTX-3::RFP neuronal marker (Supplementary Fig. 4a). While *daf-9* DA deficient mutants showed the expected low levels of *mir-84p::gfp* expression, we saw no significance difference between *dhs-26* single mutants and wild type (Supplementary Fig. 4b-c), indicating that on its own *dhs-26* does not influence DA signaling. Interestingly, however, we detected lower levels of *mir-84p::gfp* in *raga-1;dhs-26* double mutants compared to *raga-1* single mutants (Fig. 5f-g), indicating lower levels of DA signaling might emerge in the context of a genotype x genotype interaction.

Finally, to directly measure levels of DA and its precursors we turned to mass spectrometry. Consistent with our observations above, when comparing N2 to *dhs-26* single mutants, we observed no significant differences in sterol levels (Supplementary Fig. 4d). However, when comparing *raga-1;dhs-26* double mutants to *raga-1* single mutants, we found a significant reduction in Δ7-DA, (Fig. 5h), again revealing a genotype x genotype dependent interaction. Surprisingly, we also saw significantly lower levels of cholesterol and lathosterol despite unchanged pharyngeal pumping rates (Supplementary Fig. 4e), suggesting an early defect in cholesterol transport and/or metabolism in the *raga-1;dhs-26* double mutant. In addition, loss of *dhs-26* alone did not induce dauer formation (Supplementary Fig. 4f-i), or cause distal tip cell migration defects (Supplementary Fig. 4j-k), revealing that for these larval phenotypes, it does not behave like a typical DA-hormone biosynthetic gene.

We conclude that *raga-1* mutants require DHS-26 to support the uptake of sterols and production of Δ7-DA or perhaps related molecules, which acts through *daf-12* to promote longevity downstream of mTOR signaling. As a direct or indirect regulatory target of DA/DAF-12 itself, *dhs-26* presumably works in this capacity as part of a positive feedforward loop (Fig. 5i).

To gain insight into proteins and processes affected by *dhs-26* in life span regulation, we conducted single worm proteomics analyses. PCA plots revealed that mutation of *dhs-26* separates along PC2, while mutation of *raga-1* separates along PC1 (Supplementary Fig. 5a). Volcano plots showed 515 differentially down-regulated and 1093 up-regulated proteins when comparing *raga-1;dhs-26*/*raga-1* mutant animals (Supplementary Fig. 5b), from a total of 6246 detected proteins. KEGG pathway analysis revealed that down-regulated proteins were enriched for oxidative phosphorylation, lysosome biogenesis, phagocytosis, amino acid and unsaturated fatty acid biosynthesis, (Fig. 5j) as well as ER protein processing, suggesting a disruption of nutrient sensing metabolic pathways. Up-regulated proteins were enriched for RNA splicing, carbon metabolism, TCA cycle, and muscle cytoskeleton and motor protein functions, indicating altered post-transcriptional regulation and muscle remodeling (Fig. 5k). Notably, several processes show opposite regulation in *raga-1/N2* (Supplementary Fig. 2b-c) and *raga-1;dhs-26/raga-1* comparisons (Fig. 5j-k), including lysosome, beta-alanine metabolism, and nucleocytosolic transport. Collectively, these enriched pathways may indicate conflicts between nutrient sensing, catabolic and anabolic processes as responsible for the reduction of life span in *raga-1;dhs-26* animals.

Since *dhs-26* is strongly regulated by DA/*daf-12* signaling and both affect *raga-1* mutant life span, we sought to identify commonly regulated processes by cross-referencing proteomic signatures of *daf-12* and *dhs-26* mutants. In this case, we analyzed both up- and down-regulated proteins pooled together. Volcano plots comparing *raga-1;daf-12/raga-1* animals showed 102 differentially down-regulated and 112 up-regulated proteins (Supplementary Fig. 5c). When comparing proteins regulated in *raga-1;dhs-26/raga-1* to *raga-1;daf-12/raga-1* we found that 93 proteins overlapped (Supplementary Fig. 5d), which were enriched in proteins related to lysosome and muscle cytoskeleton (Supplementary Fig. 5e), suggesting that these processes may be limiting for life span.

## Discussion

### mTOR longevity operates through a bile acid-like steroid signaling pathway

Several lines of evidence indicate that DA/DAF-12 signaling functions downstream of reduced mTOR activity to promote longevity through a hormonal mechanism. Genetic epistasis analysis shows that mutations in the DA biosynthetic genes *daf-36* and *daf-9*, as well as the nuclear receptor *daf-12*, abolish the life span extension of *raga-1* mutants. Supplementation with DA restores longevity in hormone-deficient mutants but not in *daf-12* mutants, demonstrating that DA acts through its cognate receptor to promote life span extension. In contrast, mutation of the *daf-12* corepressor *din-1* does not affect *raga-1* longevity, indicating that the liganded activated receptor mediates the life span benefit.

Our results further suggest that mTOR signaling influences DA availability, consistent with a regulatory mechanism. *raga-1* mutants exhibit elevated levels of Δ7-DA despite unchanged levels of steroidogenic enzymes, implying post-translational regulation of steroid hormone production. One possibility is that reduced mTOR activity enhances the catalytic activity of biosynthetic enzymes or inhibits DA degradation, thereby increasing hormone levels. Alternatively, mTOR may regulate the uptake, trafficking, or intracellular distribution (Eid, Dauner et al. 2017) of cholesterol or DA intermediates. In contrast, disruption of steroid signaling does not measurably alter mTOR activity, supporting a model in which steroid signaling acts downstream of mTOR. Future studies will be required to identify the precise step at which mTOR influences the DA/DAF-12 pathway.

### Steroid signaling as a potential convergent mechanism of longevity

DA-dependent hormonal signaling has also been implicated in several other longevity paradigms in *C. elegans*. Life span extension in germline-deficient *glp-1* mutants (Gerisch, Rottiers et al. 2007, Shen, Wollam et al. 2012), reduced insulin signaling (*daf-2*) mutants (Dumas, Guo et al. 2013), and the dietary restriction model *eat-*2 (Thondamal, Witting et al. 2014) all depend on DA signaling, although with notable differences in pathway components. For example, Thondamal *et al*. reported that dietary restriction-induced longevity in *eat-2* mutant animals requires DA, *daf-9*/CYP27A1, and the nuclear receptor *nhr-8*/LXR, but not *daf-36* or *daf-12*. These discrepancies may reflect differences in upstream signaling inputs, sterol availability, or culture conditions. Nonetheless, the shared involvement of DA signaling across multiple independent longevity pathways suggests that bile acid-like steroid signaling represents a convergent mechanism linking diverse longevity signals to organismal physiology. The placement of steroid signaling downstream of mTOR is particularly noteworthy given the central role of mTOR in coordinating metabolic and longevity pathways.

### Conservation of the mTOR-bile acid signaling axis across species

The relationship between bile acid signaling and longevity may extend beyond nematodes. Recent work has shown that the bile acid lithocholic acid (LCA) acts as a dietary restriction mimetic. Dietary restriction in mice (Qu, Chen et al. 2024) and fasting in humans both increase endogenous LCA levels (Fiamoncini, Rist et al. 2022), and LCA supplementation extends life span in mice, *C. elegans*, and *Drosophila* (Qu, Chen et al. 2024). Although LCA has been proposed to act through the TUB-like protein 3 (TULP3) in this context, the potential contribution of bile acid-activated nuclear receptors has not been fully explored. A derivative of LCA, isoLCA is enriched in the microbiome of centenarians (Sato, Atarashi et al. 2021).

Another related derivative of LCA, 3-keto-lithocholic acid (3-keto-LCA), can activate DAF-12, the vitamin D receptor, and related bile acid–binding nuclear receptors (Motola, Cummins et al. 2006). Endogenous DAF-12 ligands, including Δ4-DA and Δ7-DA, are themselves 3-keto sterols (Mahanti, Bose et al. 2014). It is therefore plausible that metabolic conversion of LCA to 3-keto derivatives could generate ligands for nuclear receptors in this context. Furthermore, the farnesoid X receptor (FXR) agonist obeticholic acid (OCA) was shown to extend *C. elegans* life span in wild type, but not *daf-12* or *raga-1* mutants (Lijun Zhang 2025). Similarly, OCA treatment was shown to improve life span in wild type but not *Fxr-/-* mice (Lijun Zhang 2025). This suggests a broader physiological coupling between mTOR activity and bile acid signaling. Consistent with this idea, a bi-directional regulation between mTOR signaling and bile acid metabolism has been reported in several settings, including in the liver (Cai and Sewer 2013, Garcia-Rodriguez, Barbier-Torres et al. 2014, Wang, Ding et al. 2017, Wang, Chen et al. 2023).

### DHS-26 is a conserved regulator of steroid signaling and longevity

Through functional genomic screening, we identified the short-chain dehydrogenase DHS-26/DHRS1 as a key downstream effector of DA/DAF-12 signaling required for *raga-1* longevity. DHRS1 is a highly conserved NADPH-dependent reductase capable of catalyzing the reduction of steroid substrates such as estrone, androstene-3,17-dione, and cortisone *in vitro* (Wu, Xu et al. 2001, Zemanova, Navratilova et al. 2019). Despite this biochemical activity, its physiological role has remained largely unknown.

Our observations support a role for DHS-26 in mTOR-steroid signaling. mTOR inhibition leads to upregulation of *dhs-26* expression through a DA- and *daf-12*-dependent mechanism. Moreover, loss of *dhs-26*, like disruption of DA/DAF-12 signaling, abolishes life span extension in *raga-1* mutants. *raga-1;dhs-26* double mutants also display reduced levels of Δ7-DA, and supplementation with Δ7-DA restores longevity, indicating that DHS-26 influences DA production or availability. Consistent with impaired DA signaling, expression of the DAF-12 target reporter *mir-84::gfp* is reduced in the *raga-1;dhs-26* background. These observations support a model in which DHS-26 acts within the DA signaling axis to mediate mTOR-dependent life span extension.

At the same time, DHS-26 does not behave like a canonical DA biosynthetic enzyme. Mutation of *dhs-26* alone does not significantly alter DA levels; reductions in DA are observed only in the *raga-1* background, suggesting a context-dependent interaction. In addition, although *dhs-26* mutation shortens *raga-1* mutants’ adult life span, it does not produce the developmental phenotypes associated with DA deficiency, such as dauer formation or defects in gonadal migration, indicating possibly stage specific interactions. These findings suggest that DHS-26 may influence sterol metabolism more generally or produce novel ligands rather than acting at a specific step in the canonical DA biosynthetic pathway. Consistent with this possibility, *raga-1;dhs-26* double mutants reduce not only DA levels but also cholesterol levels, suggesting a broader role in regulating sterol pools, trafficking, or distribution.

Clues to its mechanism may come from our functional genomic screen and proteomic analyses, which implicated peroxisomal, lipid, and lysosomal pathways among the processes altered in *raga-1;daf-12* and *raga-1;dhs-26* mutants. Interestingly, cross-referencing our functional genomic screen with the overlap in the proteomics also revealed individual genes such as *acox-1.2,* which encodes an acyl-CoA oxidase implicated in bile acid deficiency and peroxisomal function. DHS-26 may therefore influence sterol homeostasis directly or indirectly by affecting peroxisomal or lysosomal function.

### Potential neuroendocrine roles of DHS-26

DHS-26 may also function at another scale, within neuroendocrine circuits that regulate organismal physiology. Although expressed in only a limited number of cells—including peripheral CAN neurons, a handful of head neurons, and glial cells—its loss has organism-wide effects on mTOR mediated increase of life span, consistent with a cell non-autonomous role. In particular, the prominent expression of DHS-26 in the CAN neurons raises the possibility that these cells contribute to mTOR-dependent longevity signaling. The CAN neurons are a pair of bilaterally symmetric cells with incompletely understood functions that extend processes along the excretory canal (Forrester, Perens et al. 1998). Ablation of the CANs causes larval arrest, indicating that they play essential roles in development (Forrester and Garriga 1997, Chien, Wolf et al. 2019). Interestingly, in the nematode *Pristionchus pacificus*, two genes involved in DA production, the *dauerless* gene (*dau-1*) and the hydroxysteroid dehydrogenase gene *hsd-2* (Carstensen, Villalon et al. 2021), are expressed in CAN-like cells (Mayer, Rodelsperger et al. 2015). Mutation of *dau-1* or ablation of these cells promotes dauer formation, a phenotype that can be rescued by Δ7-DA supplementation. These observations raise the possibility that CAN neurons contribute to the neuroendocrine dissemination of steroid signals in nematodes.

### Conservation of DHRS1 regulation by mTOR signaling

Consistent with a conserved endocrine role, the DHS-26 ortholog, DHRS1, shows prominent expression in several metabolically active tissues, including brain, adrenal glands, and liver (Zemanova, Navratilova et al. 2019). Notably, DHRS1 expression is also responsive to mTOR signaling. In long-lived mice treated with low-dose rapamycin, *Dhrs1* mRNA levels are elevated in the brain but unchanged in the liver (Fig. 5l) (Siegmund, Yang et al. 2017). We observe a similar inverse relationship between mTOR activity and *dhs-26* expression in *raga-1* mutants, suggesting that mTOR-dependent regulation of DHS-26/DHRS1 may be evolutionarily conserved.

Further supporting this link, mutation of the bile acid nuclear receptor FXR results in decreased hepatic expression of *DHRS1* (Fig. 5m) (Peng, Piekos et al. 2017), paralleling the regulation of *dhs-26* by DA/DAF-12 signaling in *C. elegans*. Together, these findings raise the possibility that DHS-26/DHRS1 represents a conserved node linking mTOR activity, steroid metabolism, and endocrine regulation of organismal physiology.

Future work will be required to determine whether this regulatory relationship is functionally conserved in higher organisms and to define the physiological roles of DHS-26/DHRS1 in the broader context of mTOR-regulated metabolism and aging.

## Supplements

**Supplementary Figure 1:**
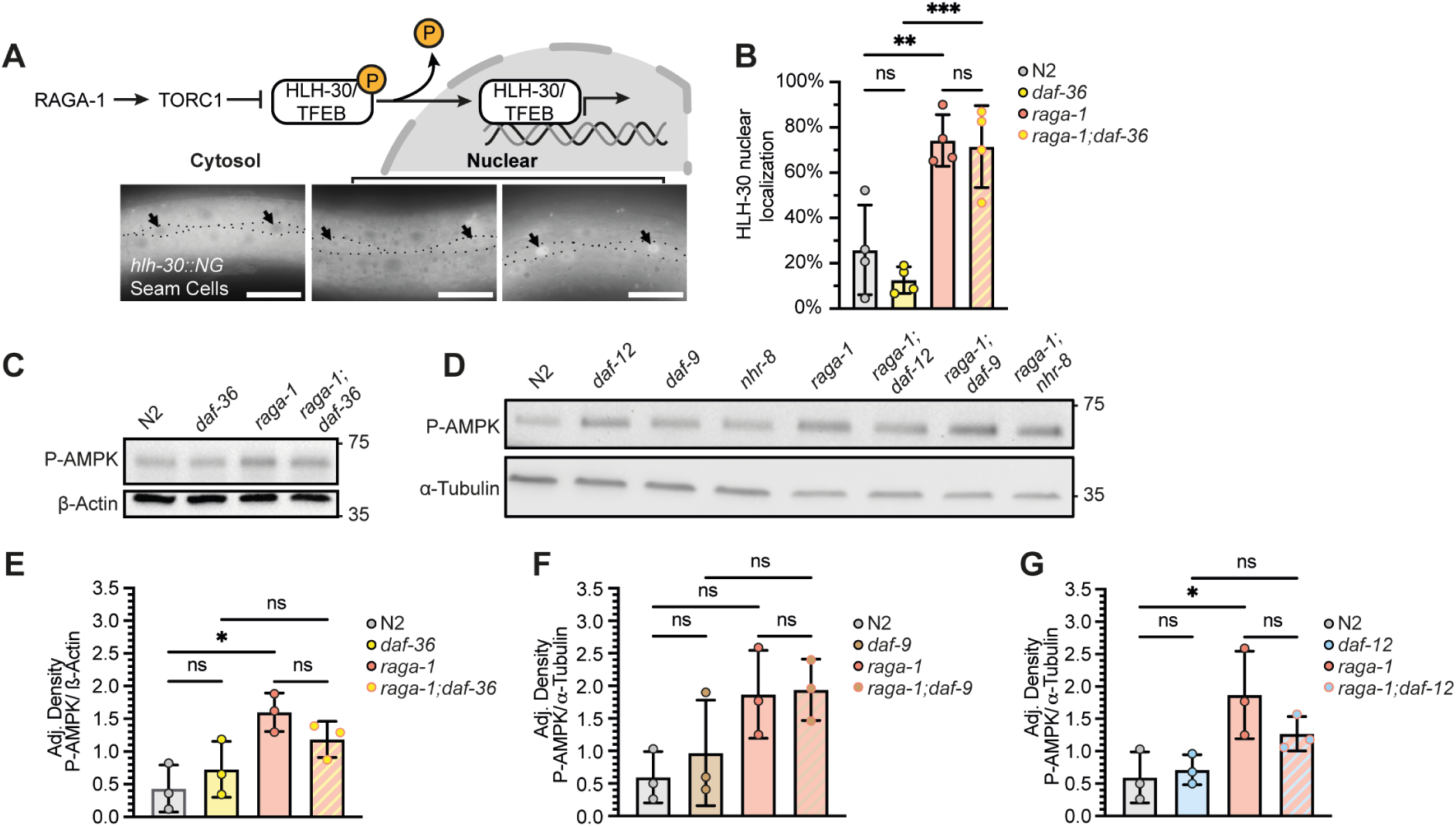
Steroid signaling does not affect mTOR activity. **a**, Schematic of TORC1 induced inhibition of HLH-30/TFEB by phosphorylation. Phosphorylation of HLH-30/TFEB inhibits its nuclear translocation and target gene expression. Cellular localization of HLH-30/TFEB was visualized and quantified using the expression of a *hlh-30::Neongreen* (HLH-30::NG) reporter in the seam cells. Representative images of N2 animals with cytosolic, and nuclear localization in seam cells, outlined and marked with arrow heads. Scale bar 20 µm. **b**, Quantification of HLH-30::NG cellular localization in seam cells at day 1 of adulthood. Statistical analysis was performed using One-way ANOVA, ns (p>0.05), ** (p<0.01), *** (p<0.005), n=4. **c-d**, Representative immunoblots with lysates from day 1 worms probed with the indicated antibodies. P-AMPK targets AMPKα phosphorylated at position Thr172. n=3. **e-g**, Quantification of immunoblots displaying P-AMPK intensities normalized to the loading controls β-Actin, or α-Tubulin. Statistical analysis was performed using One-way ANOVA, ns (p>0.05), * (p<0.01), n=3.

**Supplementary Fig. 2:**
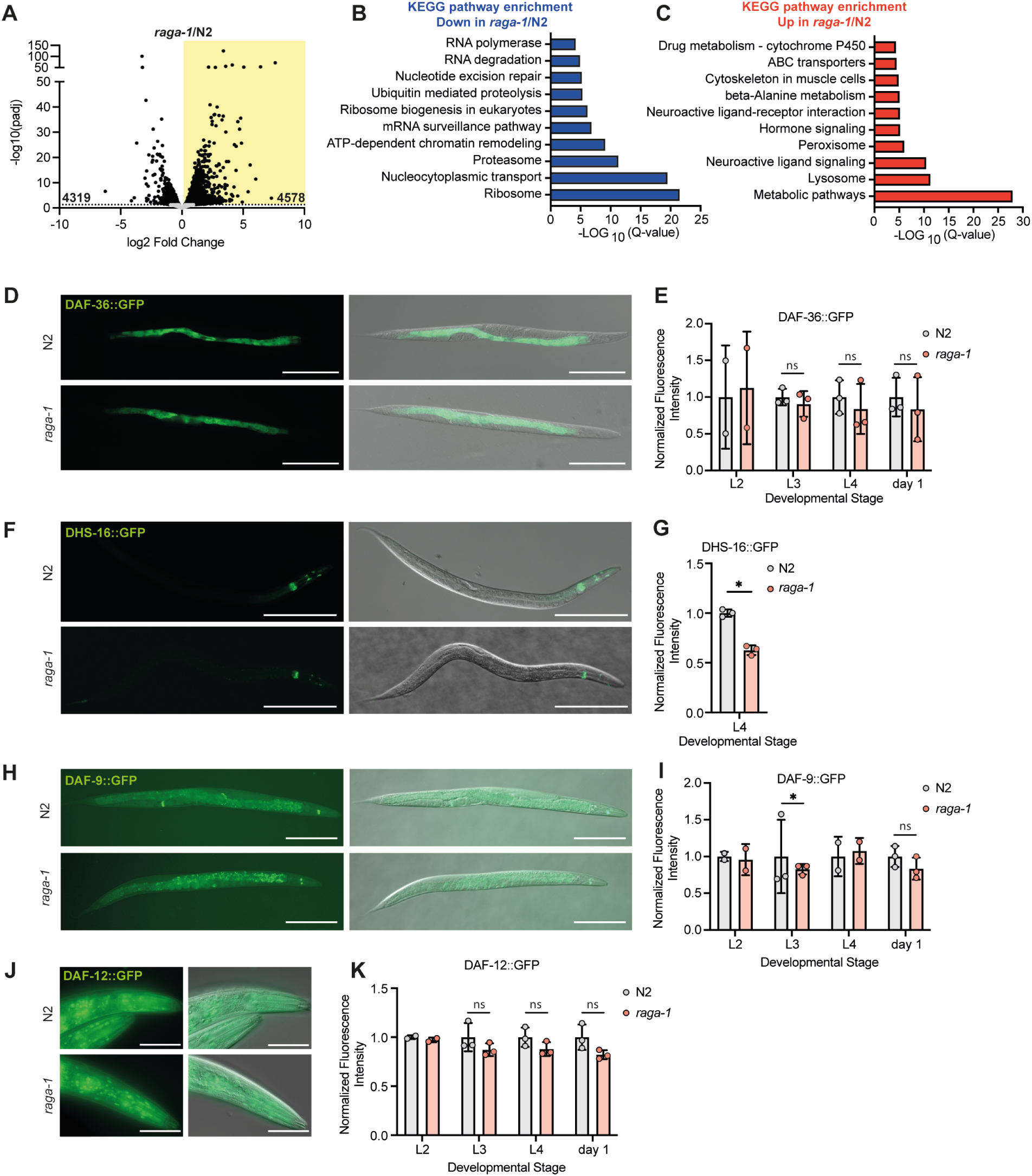
raga-1 mutation induces transcriptomic changes but does not regulate expression of steroid genes. **a**, Volcano plot of differentially expressed genes identified between *raga-1*/N2. Grey dots mark non-significantly changed genes, black dots mark significantly changed genes (Adj.p<0.05). Yellow area highlights 4578 genes significantly up-regulated, the white area shows 4319 genes that are significantly down-regulated. **b,** KEGG pathway enrichment (FDR<0.05) of significantly (Adj.p<0.05) downregulated genes (blue) comparing *raga-1(ok701)/*N2. The gene list of KEGG enrichments is listed in supplementary table 3. **c,** KEGG pathway enrichment (FDR<0.05) of significantly (Adj.p<0.05) upregulated genes (red) comparing *raga-1(ok701)/*N2. The gene list of KEGG enrichments is listed in supplementary table 3. **d**, Representative images of DAF-36::GFP expression at day 1 of adulthood comparing N2 and *raga-1(ok701)*. Differential interference contrast (DIC) and fluorescence images were taken of the animals’ full body. Scale bar 200 µm. **e**, Quantification of DAF-36::GFP expression at L2, L3, L4 and day 1 of adulthood. The developmental stage of every imaged animal was assessed by vulval morphology. **f,** Representative images of DHS-16::GFP expression at L4 stage. DIC and fluorescence images were taken of the animals’ full body. Scale bar 200 µm. **g,** Quantification of DHS-16::GFP expression at L4 stage. The developmental stage of every imaged animal was assessed by vulval morphology. **h**, Representative images of DAF-9::GFP expression at day 1 of adulthood. DIC and fluorescence images were taken of the animals’ full body. Scale bar 200 µm. **i**, Quantification of DAF-9::GFP expression at L2, L3, L4 and day 1 of adulthood. The developmental stage of every imaged animal was assessed by vulval morphology. **j**, Representative images of DAF-12::GFP expression at day 1 of adulthood. DIC and fluorescence images were taken of the animals’ head. Scale bar 50 µm. **k**, Quantification of DAF-12::GFP expression at L2, L3, L4 and day 1 of adulthood. The developmental stage of every imaged animal was assessed by vulval morphology. For e, g, i, and k, for samples with n=3 statistical analysis was performed using Student’s t-test, ns (p>0.05), * (p<0.05). Bars represent mean±SD, L2 (n=2), L3 (n=3), L4 (n=3) and day 1 of adulthood (n=3).

**Supplementary Figure 3:**
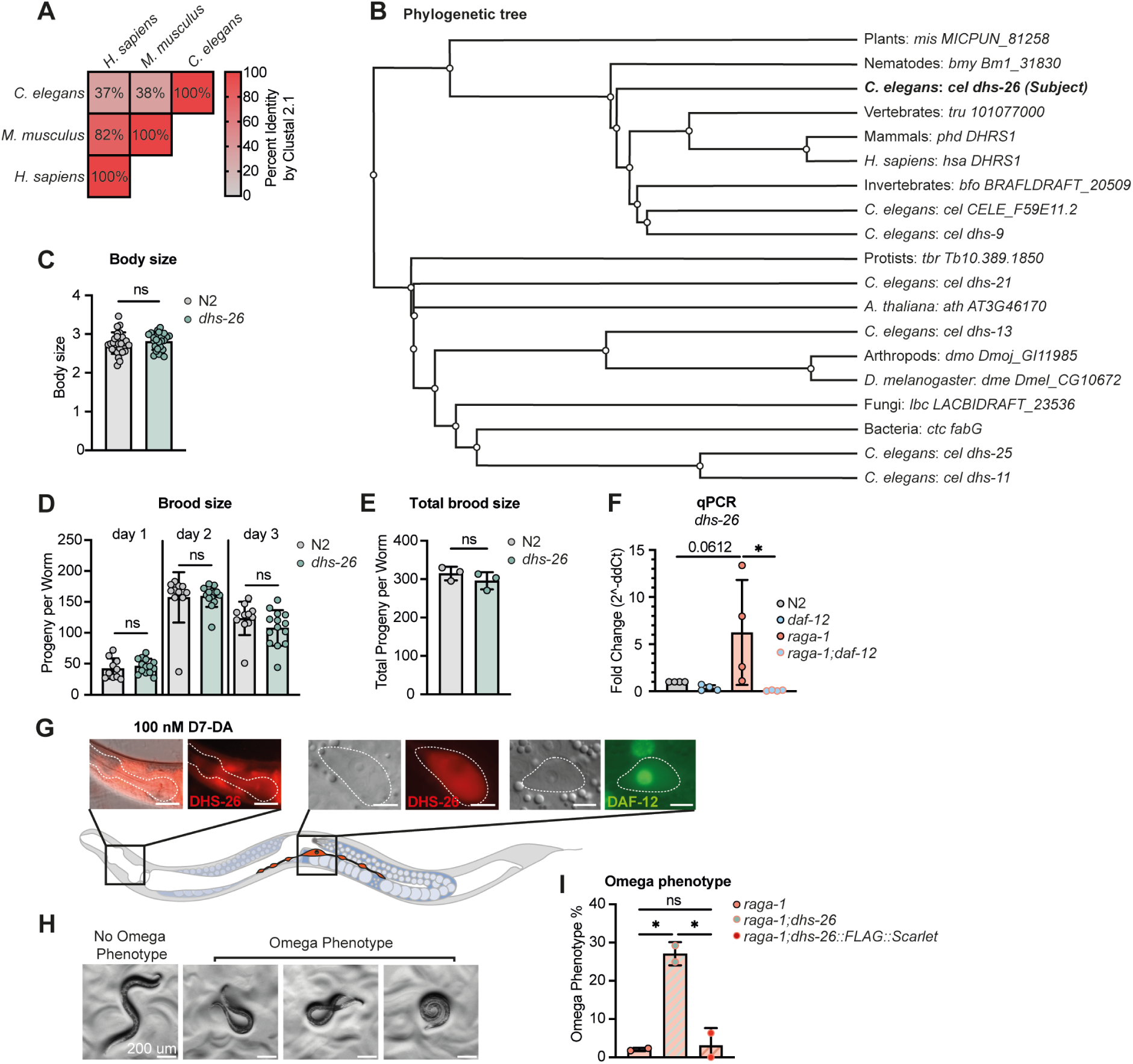
Characterization of DHS-26 conservation and dhs-26 mutation. **a**, Triangle heatmap showing percent identity between aligned DHS-26/DHRS1 proteins of different organisms using ClustalW. **b**, Phylogenetic analysis of *C. elegans dhs-26* using WormFlux. Sequence labels indicate taxonomy or model organism, organism name, and gene name, respectively. **c**, Representative body size quantified by measuring worm area comparing N2 and *dhs-26(syb8968).* Statistical analysis was performed using Student’s t-test, ns (p>0.05), n=3. **d**, Representative brood size measurement of individual N2 and *dhs-26(syb8968)* animals over a time course of 3 consecutive days starting at day 1 of adulthood. Statistical analysis was performed using Student’s t-test, ns (p>0.05), n=3. **e**, Quantification of the mean of total progeny per worm measured in (c) comparing N2 and *dhs-26(syb8968).* Bars represent mean±SD, statistical analysis was performed using Student’s t-test, ns (p>0.05), n=3. **f**, RT-qPCR quantification of *dhs-26* mRNA levels in wild type (N2), *daf-12(rh61rh411)*, *raga-1(ok701)*, and *raga-1;daf-12*. Statistical analysis was performed using One-way ANOVA, * (p<0.005), n=4. Primer sequences are listed in table 4. **g**, Representative microscopy images of DHS-26::FLAG::Scarlet (DHS-26::Scarlet) and DAF-12::GFP::FLAG of day 1 adult worms. Expression of DHS-26::FLAG::Scarlet in the head region requires supplementation of 100 nM Δ7-DA to be visible. Expression of DHS-26::FLAG::Scarlet and DAF-12::GFP::FLAG in canal associated neurons (CAN) was visible under regular growth conditions. Fluorescence and DIC images of animals’ head and mid-section. Scale bar 20 µm for the head, and 5 µm for CAN. **h**, Representative brightfield images of *raga-1;dhs-26* worms showing regular crawling (no omega phenotype) and curled body morphology (omega phenotype). Scale bar represents 200 µm. **i,** Quantification of omega phenotype penetrance comparing *raga-1(ok701)*, *raga-1;dhs-26*, and *raga-1;dhs-26::FLAG::Scarlet*. Bars represent mean±SD, statistical analysis was performed using Student’s t-test, ns (p>0.05), * (p<0.05), n=2.

**Supplementary Figure 4:**
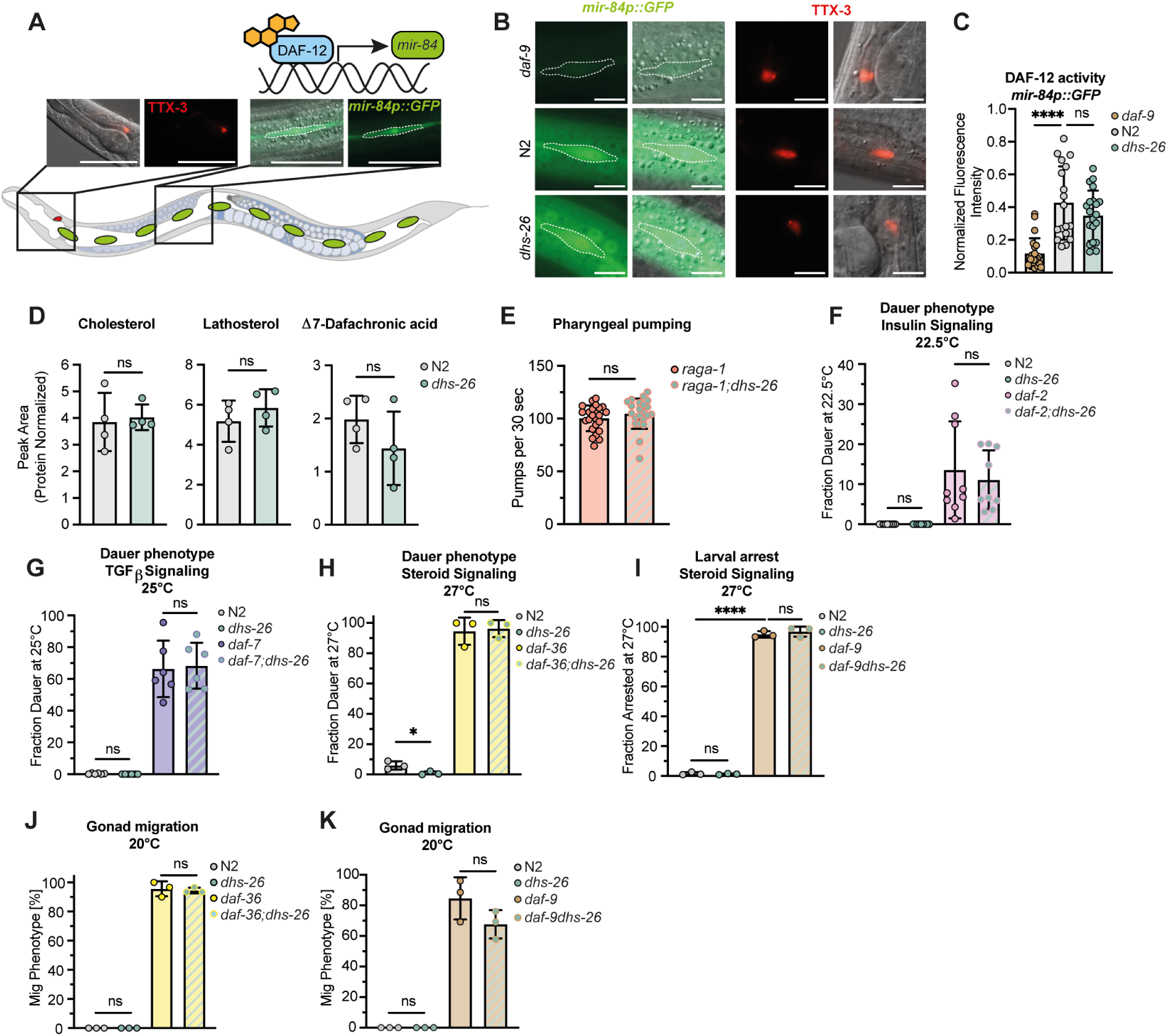
DHS-26 does not function as a canonical DA biosynthetic enzyme in wild type animals. **a**, Schematic of the positive regulatory arm of Δ7-DA-bound DAF-12 resulting in the transcriptional activation of its target gene *mir-84*. Representative images of the *mir-84p::GFP* expression pattern in seam cell V2 at larval stages L4.3-L4.5 (Mok, Sternberg et al. 2015). The larval stage of every imaged animal was assessed by vulval morphology. Differential interference contrast (DIC) and fluorescence images were taken of the midbody for *mir-84p::GFP*, and head for TTX-3::RFP. Scale bar represents 50 µm. **b**, Representative images of *mir-84p::GFP* in seam cell V2 of animals at larval stages L4.3-L4.5 (Mok, Sternberg et al. 2015). The larval stage of every imaged animal was assessed by vulval morphology. DIC and fluorescence images were taken of the animals’ midbody for *mir-84p::GFP*, and head for TTX-3::RFP. Scale bar represents 10 µm. **c**, Representative quantification of *mir-84p::GFP* expression in seam cell V2 of N2, *dhs-26(syb8968)*, and *daf-9(rh50)* animals. Fluorescence intensity of each region of interest (ROI) was calculated by subtracting the background fluorescence of each image from the ROI. *mir-84p::GFP* was normalized to TTX-3::RFP expression in AIY interneurons of the same animal. Statistical analysis was performed using One-way ANOVA, ns (p>0.05), **** (p<0.001), n=4. **d**, Relative concentrations of Δ7-dafachronic acid (DA) and its precursors cholesterol and lathosterol extracted from wild type (N2) and *dhs-26(syb8968)* day 1 adults. Measured concentrations were normalized to protein levels. Statistical analysis was performed using Student’s t-test, ns (p>0.05), n=4. **e**, Representative measurement of pharyngeal pumping rates of day 1 adults measured in 30 s. Bars represent mean±SD, statistical analysis was performed using Student’s t-test, ns (p>0.05), n=3. **f-i**, Quantification of dauer phenotype penetrance 48h after egglay, assessing mutation in *dhs-26* on the following pathways: insulin signaling at 22.5°C (e, n=9), TGFβ signaling at 25°C (f, n=6), and steroid signaling in *daf-36(k114)* (g, n=3) and *daf-9(rh50)* (h, n=3) at 27°C. Bars represent mean±SD, statistical analysis was performed using One-way ANOVA, ns (p>0.05), * (p<0.05), **** (p<0.0001). **j-k**, Quantification of *dhs-26(syb8968)* effects on distal tip cell migration defects in L4 animals grown on cholesterol-free plates egg-on in steroid signaling mutants *daf-36(k114)* (i) and *daf-9(rh50)* (j). Bars represent mean±SD, statistical analysis was performed using One-way ANOVA, ns (p>0.05), n=3.

**Supplementary Figure 5:**
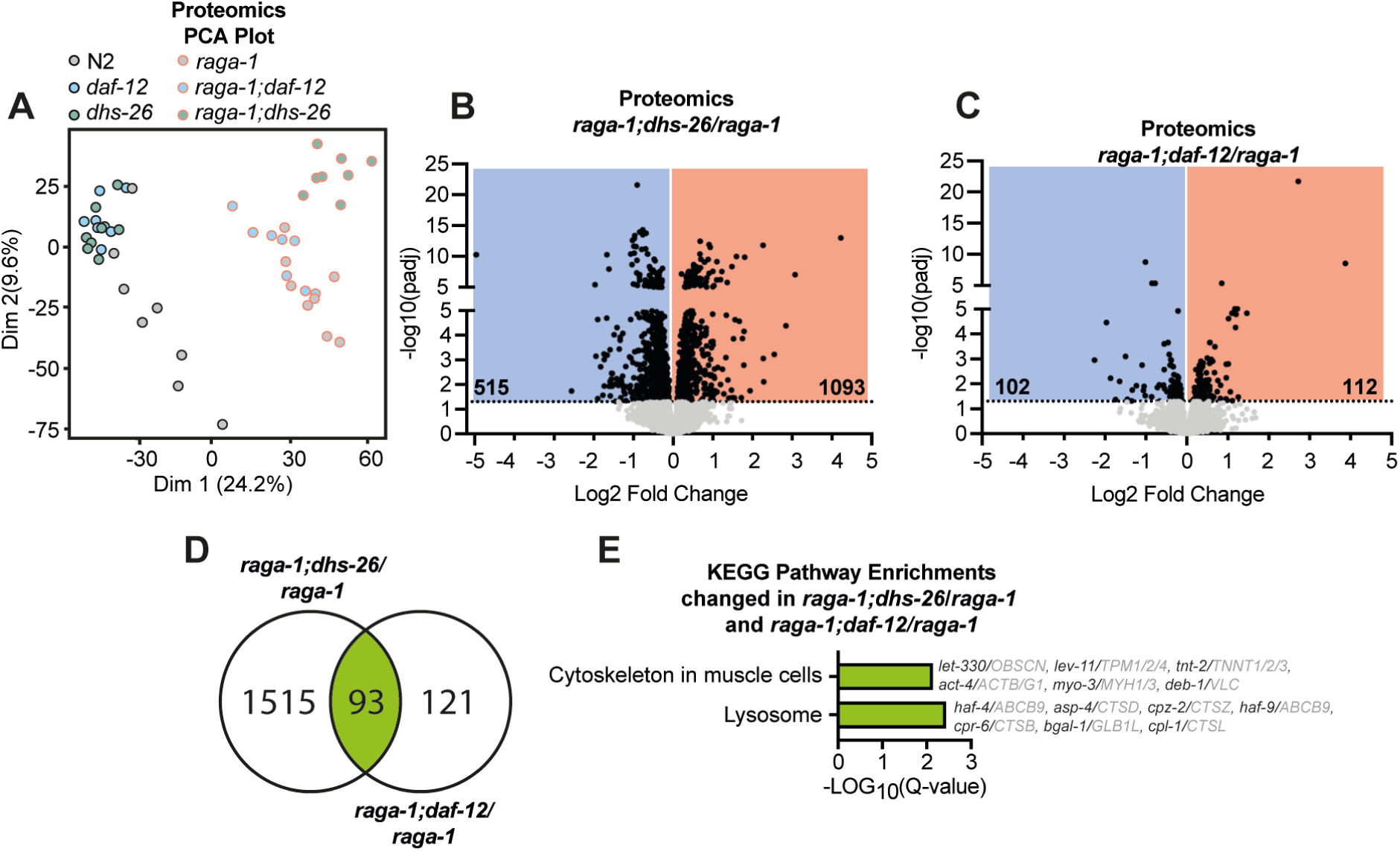
Proteomics changes induced by loss of dhs-26 overlap with changes induced by mutation of daf-12. **a**, PCA plot of single worm proteomics data comparing wild type (N2), *daf-12(rh61rh411)*, *dhs-26(syb8968)*, *raga-1(ok701)*, *raga-1;daf-12*, and *raga-1;dhs-26* in day 1 adults. **b**, Volcano plot of differentially expressed genes comparing *raga-1;dhs-26/raga-1(ok701).* Black dots mark significantly changed genes (Adj.p<0.05), grey dots mark non-significantly changed genes (Adj.p>0.05). Blue area highlights 515 significantly down-regulated genes, the red area highlights 1093 significantly up-regulated genes. **c**, Volcano plot of differentially expressed genes comparing *raga-1;daf-12/raga-1(ok701).* Black dots mark significantly changed genes (Adj.p<0.05), grey dots mark non-significantly changed genes (Adj.p>0.05). Blue area highlights 102 significantly down-regulated genes, the red area highlights 112 significantly up-regulated genes. **d**, Venn diagram overlapping significantly regulated proteins comparing *raga-1;dhs-26/raga-1(ok701)* and *raga-1;daf-12/raga-1(ok701),* resulting in the identification of 93 shared genes. **e**, KEGG pathway enrichment (FDR<0.05) of the 93 shared genes shown in f. Gene list and the respective ortholog of the enriched pathways is displayed. The gene list of KEGG enrichments is additionally listed in supplementary Table 3.

## Acknowledgements

We would like to thank the MPI-AGE core facilities for their services, including the Bioinformatics, Proteomics and Metabolomics cores, members of the Antebi lab for helpful discussions, and Sarah Kreuz and Stephanie Fernandez for critical reading of the manuscript. Klara Schilling was supported by the International Max Planck Research School, Cologne Graduate School of Ageing Research and the project funded through the Max Planck Gesellschaft. The Austrian Science Fund FWF, DK-MCD W1226 to A.Z.

## Author Contributions

A.A and K.S. conceived and designed the study. A.A., A.Z., and T.M. laid the foundation to this study. Investigation, K.S., K.M.M. (DHS-16::GFP microscopy), and A.L. (support with life span screen). R.S. and H.J.K. synthesis and providing of Δ7-DA. Writing, review and editing, A.A., and K.S.

## Methods

### Materials availability

Information and requests for resources and reagents should be directed to the corresponding author. This study did not generate unique reagents or new plasmids. Strains generated in this study are available upon request.

### Experimental model and subject details

Nematodes were cultured using standard techniques at 20 °C on NGM agar plates containing 5 µM Cholesterol, seeded with the *Escherichia coli* strain OP50, unless otherwise noted. All strains used in this study are listed in the table 4. Mutant strains were obtained from the Caenorhabditis Genetics Center (CGC) and outcrossed to our N2 WT (*C. elegans* variant Bristol) as indicated in table 4. CRISPR–Cas9 mutants were generated by SunyBiotech (https://www.sunybiotech.com/).

### Life span experiments

All life spans were performed at 20 °C. For each genotype, ∼150 adults worms (6 x 6 cm plates with 30 worms each) were grown on standard NGM plates with OP50. Survival was monitored every other day. Worms that did not respond to gentle touch by a worm pick were scored as dead and were removed from the plates. Animals that crawled off the plate or had ruptured vulva phenotypes were censored. All life span experiments were blinded. Graphpad Prism (9.0.0) was used to plot survival curves. Survival curves were compared and *p*-values were calculated using the log-rank (Mantel-Cox) test. Complete life span data are available in supplementary Table 1.

### RNAi in C. elegans

RNAi experiments were performed as previously described. *E. coli* HT115 were used and animals were fed with the desired RNAi bacteria from egg on. One exception being RNAi knockdown of *let-363* and its *luci* control for which eggs were grown on NGM plates seeded with OP50 bacteria and worms were transferred to *E. coli* iOP50 bacteria on day 1 of adulthood. The HT115 bacteria were from the Vidal or Ahringer library. The iOP50 competent bacteria were transformed with double-stranded RNA expressing plasmids, which were extracted from the respective HT115 bacterial strains. RNAi bacteria were grown in lysogeny broth medium supplemented with 100 µg ml−1 ampicillin at 37 °C overnight with gentle shaking. The culture was spread on RNAi plates, which are NGM plates containing 100 µg ml−1 ampicillin and 0.4 mM isopropyl β-D-1-thiogalactopyranoside. RNAi-expressing bacteria were allowed to grow on the plates at room temperature for 2 days.

### Functional genomics screen

Worms were synchronized by egg lay of day 1 adults on a NGM OP50-seeded plates. Eggs were manually transferred to RNAi-seeded plates of the respective candidate RNAi. RNAi plates were prepared as described above. L4 worms were again transferred to fresh RNAi plates for a total of around 150 worms per condition (6 x 6 cm plates with 30 worms each). Life span was conducted as described above, scoring for survival every 2-3 days. For controls *raga-1(ok701)* and *raga-1;daf-12(rh61rh411)* grown on luciferase RNAi were included in every life span experiment.

### Dauer assay

To determine dauer formation, synchronized worm populations were generated by allowing 20 worms to lay eggs for 4 h. Eggs were incubated at 22.5°C, 25 °C, or 27°C respectively for 48 h. Dauer characteristics such as a dark coloration and a long, thin body shape were used for identification (Riddle and Albert 1997). The development of a minimum of 100 worms was scored fort each condition.

### Defective gonad migration (Mig) assay

To determine gonad migration defects, synchronized worm populations were generated by allowing 20 worms to lay eggs for 4 h. Eggs were incubated at 20°C, or 25 °C respectively for 48 h. Animals were scored as Mig (Hedgecock, Culotti et al. 1987) if one of the two gonad arms failed to turn. At least 20 worms were analyzed per genotype.

### Omega morphology assay

To determine the prevalence of a body curling (omega) phenotype, day 1 worms were allowed to lay eggs for 4 h. Eggs were incubated at 20°C for 72 h. Moving animals were scored as omega if there was contact between anterior and posterior (Supplementary Fig. 3h). At least 20 worms were analyzed per genotype.

### Pharyngeal pumping assay

To assess the pharyngeal pumping rate, L4 animals were picked and allowed to develop to day 1 adults at 20 °C over night. Pharyngeal pumping was counted for 30 seconds. 20-25 worms were analyzed per condition.

### Worm size measurements

Images of worms were taken with a Zeiss Axio Imager Z1 or a Leica M165 FC microscope. Body length was determined using ImageJ. At least 25 worms were analyzed per genotype.

### Brood size measurements

To determine brood size, at least 10 L4 animals were isolated for each genotype. Animals were transferred to fresh plates after 24h, 48h, and 72h and brood size was counted for each day.

### RNA extraction and qPCR analysis

Worms were synchronized by egg lay of day 1 adults and were collected at early day 1 stage. Worms were collected in 15 mL M9 buffer and washed 3x with M9 to remove bacterial contaminations. Worms were then resuspended in 700 µl QIAzol lysis reagent (Qiagen), snap-frozen in liquid nitrogen, and stored at -80°C until extraction. To lyse the worms, 200 µL of 0.1 mm Zirconia/Silica beads (FisherScientific) were added to the thawed worm suspension and transferred to the TissueLyser LT (Qiagen) for 15 min, full speed at 4°C. RNA extraction and purification were performed using the RNeasy Mini Kit (Qiagen) and DNase digestion was carried out using the RNase-Free DNase Set kit (QIAGEN) according to manufacturer’s instructions. RNA was eluted in 50 µl RNase-free water. RNA purity and concentration were assessed using the NanoPhotometer NP80 spectrophotometer (Implen). RNA was stored at - 80°C. cDNA synthesis was carried out using iScript (Bio-Rad). Quantitative PCR was performed using Power SYBR Green Master Mix (Applied Biosystems) on a ViiA 7 Real-Time PCR System (Applied Biosystems), using 40 cycles of 95°C for 15 s and 60°C for 1 min with a temperature change of 1.6°C/s. Four technical replicates were averaged for each sample and primer pair. Before the experiment, all primer sets were validated using standard dilution curves. *act-1* and *cdc-42* were used as internal references, see table 4 for primer sequences.

### RNA-seq C. elegans

RNA was extracted as described above. The RNA libraries were prepared by the Cologne Center for Genomics (CCG). The removal of ribosomal RNA was conducted using the RiboZero rRNA removal kit (Illumina). Sequencing was performed on Illumina HiSeq4000 sequencing system (25 million reads per sample) using a paired-end 2 x 100-nt sequencing protocol. Sequencing analysis was performed by the MPI bioinformatics core facility. In brief, Ensembl *Caenorhabditis elegans* release 105/ce11 was used as the reference sequence, rRNA transcripts were removed from the annotation file and reads were pseudo aligned to the reference transcriptome and were quantified using kallisto/0.46.1 (Bray, Pimentel et al. 2016). Read counts were normalized by making use of the standard median-ratio for estimation of size factors and pair-wise differential gene expression was calculated using DEAes/1.24.0 (Love, Huber et al. 2014). Genes with fewer than ten overall reads were removed and overall log2 changes were shrunk using approximate posterior estimation for GLM coefficients. Transcriptomics data is available in Supplementary Table 5.

The KEGG pathway analysis of significant genes was performed using the DAVID bioinformatics resource database. Statistical analysis was performed using the Benjamini and Hochberg procedure (FDR), FDR<0.05 for significant enrichment.

### Single worm proteomic sample preparation and analysis

Single young adult worms were placed into pre-chilled 0.1 mL PCR strip tubes containing 4 μL lysis buffer (0.25% DDM, 125 mM TEAB) and snap-frozen in liquid nitrogen within 3 min. 8 single worms were used for each condition. Samples were stored at −80°C until processing. For lysis, worms underwent four freeze–thaw cycles (liquid nitrogen; water at room temperature) followed by sonication in a Bioruptor (Diagenode S.A.) coupled to a Minichiller 300 (Huber) (20 cycles, 30 seconds on, 30 seconds off) at 4°C. Digestion was initiated by adding 5 μL chilled PBS pH 7.4 and 1 µL of a trypsin/Lys-C mix (100 ng/μl Trypsin and 50 ng/μl LysC in 5 mM acetic acid). Samples were then incubated at 37°C o/n in a thermocycler. Digests were loaded onto EvoTips for LC–MS/MS.

The Evosep One liquid chromatography system (Bache, Geyer et al. 2018) was used for analyzing the samples with the predefined 30 samples per day (30SPD) method. The analytical column used was an ReproSil-Pur column, 15 cm x 150 μm, with 1.9 μm C18 beads (EV1106 Endurance Column, Evosep). The mobile phases A and B were 0.1 % formic acid in water and 0.1% formic acid in 100% ACN, respectively.

Peptides were analyzed on a hybrid trapped ion mobility spectrometry (TIMS) quadrupole time-of-flight (TOF) mass spectrometer (timsTOF Pro 2, Bruker) in a data-independent acquisition parallel accumulation, serial fragmentation (diaPASEF) mode. The mass spectra range was set to 100-1700 m/z and TIMS ion accumulation and ramp times were set to 100 ms and total cycle time was 2.0 s. The ion mobility range was set to 1/K0 = 0.8-1.25 V s/cm^2^. Isolation windows in the m/z versus ion mobility plane were defined to cover the region of highest precursor ion density with an m/z slice width of 26 Th. Collision energy was applied linearly with ion mobility from 0.6 to 2.0 V s/cm^2^, and collision energy from 20 to 59 eV.

Raw data were analyzed using Spectronaut version 19.3.24 (Biognosys) using the default parameters against the one-protein-per-gene reference proteome for *C.elegans*, UP000001940, downloaded August, 2022. Methionine oxidation and protein N-terminal acetylation were set as variable modifications; cysteine carbamidomethylation was set as fixed modification. The digestion parameters were set to “specific” and “Trypsin/P,” with two missed cleavages permitted. Protein groups were filtered for at least two valid values in at least one comparison group and missing values were imputed from a normal distribution with a down-shift of 1.8 and standard deviation of 0.3. Differential expression analysis was performed using limma, version 3.60.6 (Ritchie, Phipson et al. 2015), in R, version 4.4.0. Proteomics data is available in Supplementary Table 6.

### Targeted Metabolomics sample preparation and analysis

Worms were synchronized by egg lay of day 1 adults and were grown on NGM plates supplemented with cholesterol of high purity (Sigma, ≥99%). 10,000 staged day 1 worms were collected in 15 mL M9 buffer and washed 3x with M9 to remove bacterial contaminations. Worm pellets were harvested, washed and snap-frozen in liquid nitrogen within 15 min. Samples were stored at -80 °C until extraction.

Samples were taken from -80°C freezer and stored on dry ice. Sequentially, 200 μl zirconium beads (1mm) and 1 ml of extraction buffer (50 ml MeOH, 20 μl Cholesterol D7) were added and samples were immediately lysed in a tissue lyser for 30 min at 50 Hz at 4°C. Samples were transferred to 5 ml tubes and 3 ml of methanol were added. To fully homogenized worms, the samples were further sonicated with a Bioruptor Plus (Diagenode S.A.) coupled to a Minichiller 300 (Huber) (20 cycles, 30 seconds on, 30 seconds off) in a water bath filled with ice and with the zirconium beads still in the sample. From the total volume of 4 ml, an aliquot of 200 μl from the homogenized and mixed samples was transferred to a fresh 1.5 ml Eppendorf tube and centrifuged at 21,000 rpm at 4°C for 10 min to obtain a protein pellet for quantification. To get a cleared extract, 5 ml tubes were centrifuged at 4°C at 21,000 rpm for 10 min. 1700 µl supernatant were used for LC-MS analysis and 1700 μl for GC-MS analysis. All samples were dried in a speed vac (ScanSpeed40) and stored at -80 °C until analysis.

Cholesterol and Lathosterol were measured using GC-MS (Gas Chromatography coupled to a Q-Exactive-Orbitrap mass spectrometer, Thermo Fisher Scientific). For this purpose, metabolites were derivatized using a two-step procedure starting with a methoxyamination (methoxyamine hydrochlorid, Sigma) followed by a trimethyl-silylation using N-Methyl-N-trimethylsilyl-trifluoracetamid (MSTFA, Macherey-Nagel). Dried samples were methoxyaminated by re-suspending them in 10 μL of a freshly prepared methoxyamine in pyridine (Sigma) solution (40 mg/mL). The samples were incubated for 45 min at 40 °C on an orbital shaker (VWR) at 1500 rpm. In the second step 90 μL of MSTFA spiked with C8 - C40 Alkane standard (40147-U, Sigma Aldrich) to a concentration of 1 μg/ml was added and the samples were incubated for additional 45 min at 40°C and 1500 rpm. At the end of the derivatization the samples were centrifuged for 2 min at 21,100x g and the clear supernatant was transferred to fresh auto sampler vials with conical glass inserts (300 μl, Chromatographie Zubehoer Trott).

For the GC-MS analysis 0.5 μL of each sample were injected using a TriPlus RSH autosampler system (Thermo Fisher Scientifc) using a Split/SplitLess (SSL) injector at 250°C in splitless mode. The carrier gas flow (helium) was set to 1 ml/min using a 30m MEGA-5 MS capillary column (0.250 mm diameter and 0.25 μm film thickness, MEGA). The GC temperature program was: 1 min at 70°C, followed by a 9 °C per min ramp to 350 °C. At the end of the gradient the temperature is held for additional 5 min at 350 °C. The transfer line and source temperature are both set to 280°C. The filament, which was operating at 70 eV, was switched on 4.5 min after the sample was injected. During the whole gradient period the MS was operated in full scan mode covering a mass range m/z 70 and 700 with a resolution of 60,000. The GC-MS data analysis was performed using the open-source software El Maven (Agrawal, Kumar et al. 2019) (Version 0.12.0). For this purpose, Thermo raw mass spectra files were converted to mzML format using MSConvert (Chambers, Maclean et al. 2012) (Version 3.0.22060, Proteowizard). The identity of each compound was validated by authentic reference compounds, which were measured at the beginning or at the end of the sequence, furthermore, a compound’s identity was matched to EI spectra and the retention index (RI).

For data analysis the peak areas of extracted ion chromatograms from selected fragment ions were determined with El Maven; only in rare cases peaks were manually re-integrated. Extracted ion chromatograms were generated with a mass accuracy of <5 ppm and a retention time (RT) tolerance of <0.05 min as compared to the independently measured reference compounds. These areas were then normalized to the internal standards which were added to the extraction buffer followed by a normalization to the protein content of the analyzed samples.

Δ7-Dafachronic acid was measured using LC-MS. Dried samples were reconstituted in 100 μl 80% MeOH for 10 min on a shaker with 1500 rpm at 4°C. Samples were centrifuged at 21,000 rpm for 10 min at 4°C and supernatant was transferred to glass LC Autosampler Vials with 300 μl glass inserts. 20 µl of the sample were injected and analyzed in a Vanquish horizon UHPLC coupled to a QExactive plus HRMS (Thermo Fisher Scientific).

Separation on the UHPLC system was done on a HSS T3, 2.1 x 150 mm column with precolumn and a gradient elution with buffer A (85% MeOH) and buffer B (100 % MeOH) both with 5 mM Ammonium acetate. The column was kept at 30 °C and the flow rate was 300 μl/min and the gradient as follows: 0-2 min, 0% B; 2-15 min 0-100% B, 15-25 min, 100%B and 25.1-27.5, 0% B. The MS was running in negative mode from 0-15 min and in positive mode from 15-27.5 min. Δ7-Dafachronic acid was analyzed with a t-SIM scan and identity was validated by running pure reference compounds. Data was extracted using TraceFinder (v. 5.1, Thermo Fisher Scientific) software by integrating the respective peak in the t-SIM scans of the [M-H]-and [M+H]+ adduct mass traces with a mass accuracy of <5 ppm and a retention time (RT) tolerance of <0.05 min as compared to the independently measured reference compounds.

### Imaging and image analysis

For live imaging, worms were anesthetized in 0.1 % sodium azide and imaged using a Zeiss Axio Imager Z1 microscope. Image analysis was performed using Fiji software (Schindelin, Arganda-Carreras et al. 2012). For *hlh-30::mNeonGreen* nuclear localization, L4 worms were anesthetized for 60 seconds and scored in the seam cells under blinded conditions for cytosolic and nuclear localization for a maximum of 3 minutes. Gene expression of *daf-36::GFP*, *daf-9::GFP*, and *daf-12::GFP* was scored at different stages of development (L2, L3, L4, and day 1 of adulthood). For *daf-36::GFP*, *dhs-16::GFP*, and *daf-9::GFP* expression in the full body was measured. For *daf-12::GFP* expression in the head neurons was quantified. *dhs-26::mScarlet* expression was quantified in the CAN of day 1 adult animals. *dhs-26::mScarlet* expression was imaged in the head of day 1 adult animals. *daf-12::GFP::FLAG* expression was imaged in the CAN of day 1 adult animals. *mir-84::GFP* expression was determined in the seam cells of L4 larvae and was normalized to *ttx-3::RFP* expression in the ASI head neurons. The fluorescence intensity was determined by defining regions of interest (ROIs) and subtracting the background fluorescence from the ROI. This was repeated for each taken image.

### Δ7-Dafachronic acid supplementation

NGM plates were prepared and seeded with OP50 *E. coli* bacteria. Ready to use plates were freshly infused with steroids the day of use by pipetting 50 µl Δ7-Dafachronic acid in 100% EtOH on top of the bacterial lawn of a 6 cm plate. For a final concentration of 100 nM, a stock solution of 20 µM was used. For a final concentration of 500 nM, a stock solution of 100 µM was used. Control plates were infused with 50 µl EtOH vehicle only. The concentration of the used steroid stock solution was determined by calculating the final concentration in relation to total agar volume.

### Western blotting

Worms were synchronized by egg lay of day 1 adults and were collected at day 1 adult stage. Worms were collected in 15 mL M9 buffer and washed 3x with M9 to remove bacterial contamination. Samples were then snap-frozen in liquid nitrogen and stored at -80°C until lysis. The pellet was resuspended in 100 µl lysis buffer supplemented with 1 unit for 10 ml cOmplete ULTRA EDTA-free protease inhibitors (Roche) and 1 unit for 10 ml PhosSTOP phosphatase inhibitors (Roche) and was sonicated with Bioruptor Plus (Diagenode S.A.) coupled to a Minichiller 300 (Huber) for 15 cycles of 30 sec sonication, 30 sec rest at 4 °C. The total protein extract was cleared by spinning down at 13,000x g for 10 min at 4 °C. Supernatant protein was quantified using Pierce BCA Protein Assay Kit (ThermoFisher Scientific) according to the manufacturer’s instructions. Reagents and samples were incubated at 37 °C for 30 mins before measurement. Equal amounts of protein were mixed with 4X Laemmli buffer with 0.9% 2-mercaptoethanol and incubated at 95°C for 5 min, and then stored at -20 °C until use.

Before SDS-PAGE, samples were again incubated at 95°C for 5 min and protein was then loaded on a midi protein gel (Bio-Rad). For P-AMPK/β-Actin in *daf-36* mutant animals 30 µg protein per lane were loaded. For P-AMPK/α-Tubulin in *daf-9, daf-12, and nhr-8* mutant animals 40 µg protein per lane were loaded. For FLAG/β-Actin in *raga-1* mutant animals 75-100 µg protein per lane were loaded. The gel was blotted to a midi nitrocellulose membrane (Bio-Rad) using the Trans-Blot Turbo Transfer System (Bio-Rad). The membrane was blocked for 1 h in 5 % skim milk in tris-buffered saline with Tween20 (TBST) (24.77 mM Tris base, 137 mM NaCl, 2.68 mM KCl, 0.05% Tween20, pH 7.4) and incubated with primary antibody overnight at 4 °C. Primary antibodies were diluted as follows: anti FLAG 1:500 in 5 % BSA, anti β-Actin 1:1000 in 5 % BSA, anti α-Tubulin 1:2000 in 5 % skim milk, anti P-AMPK(Ser211) 1:1000 in 5 % skim milk. Next, the membrane was incubated with the secondary antibody for 2 h at room temperature using dilutions of 1:10,000 in 5 % BSA for anti mouse and anti rabbit. Membrane was imaged using Western Lightning Plus Enhanced Chemiluminescence Substrate (PerkinElmer). Used antibodies are listed in table 4.

### Statistics and reproducibility

The statistical tests performed in this study are indicated in figure legends and method details. If not stated otherwise in the figure legends, data are represented as the mean ± s.d.

Number of replicates and animals for each experiment are enclosed in their respective figure legends or method details. Data collection and analysis were performed blind to the conditions of the experiments.

For normally distributed data, unpaired *t*-test with Welch’s correction or one-way ANOVA with Brown-Forsythe and Welch’s correction were calculated in GraphPad Prism 9 (GraphPad software). For life span analysis, *p*-values were calculated using log-rank (Mantel-Cox) test. Statistical data for life spans (except for RNAi screen life spans) can be found in supplementary table 1. Statistical data from RNAi screen life spans can be found in supplementary table 2. Genes included in KEGG enrichment terms are listed in supplementary table 3. Significance levels are: not significant (ns) *p*>0.05, * *p*<0.05. ** *p*<0.01, *** *p*<0.001, **** *p*<0.0001.

### Data availability

All data generated in this study are available in the main text or the Supplementary Information. *C. elegans* RNA-seq data accession code: XXXXX; WBcel235.80 reference genome *C. elegans* RNA-seq. *C. elegans* Proteomics data accession code: XXXXX; UP000001940_6239 reference proteome *C. elegans*.

## Additional Files

Supplementary table 1: Lifespan tables

Supplementary table 2: RNAi screen lifespan table

Supplementary table 3: KEGG enrichment gene list

Supplementary table 4: Reagents and resources list

Supplementary table 5: Transcriptomics master table

Supplementary table 6: Proteomics master table

9 Source data tables: Fig. 1-5, Supplementary Fig. 1-4

## Abbreviations

DA: Dafachronic acid
FXR: farnesoid X receptor
mTOR: mechanistic target of rapamycin
NHR: nuclear hormone receptors
VDR: vitamin D receptor
DEGs: differentially expressed genes
luc: luciferase
COBALT: constraint-based multiple sequence alignment tool
CAN: canal associated neuron
ROI: region of interest
LCA: lithocholic acid
TULP3: TUB-like protein 3
3-keto-LCA: 3-keto-lithocholic acid
OCA: obeticholic acid

